# Tripeptidyl peptidase II is essential for maintaining cerebrovascular homeostasis of female mice and represents a novel therapeutic target for vascular dementia

**DOI:** 10.64898/2026.05.05.722100

**Authors:** Jin Zhao, Qing Guo, Xiaohong Xin, Yujie Zhang, Kaixuan Pei, Junyao Li, Gang Wang, Jinghui Li, Wanqi Sun, Sai Huang, Xueyu Fan, Linlin Luo, Wen Li, Yuqi Wang, Yuanyuan Cao, Bin Yang, Hairong Zhang, Shichang Yang, Ruiling Zhang, Jie Liu, Junqiang Zhao, Xiaochuan Zhang, Jisen Huai

**Author notes:** Co-senior author for imaging, anatomy, and related analysis (Jie Liu, Junqiang Zhao, Xiaochuan Zhang). correspondence (J.H.). These authors contributed equally (Jin Zhao, Qing Guo, Xiaohong Xin).

## Abstract

Although cerebrovascular impairment is a known common driver of both Vascular dementia (VaD) and Alzheimer’s disease (AD), the underlying mechanisms remain poorly defined. Substantial evidence has demonstrated that brain-derived estrogen is essential for cerebrovascular health and neuroprotection. Consequently, therapeutic strategies that can replicate the beneficial effects of estrogen replacement therapy (ERT) in the brain while avoiding its peripheral risks are being actively pursued. Here, we demonstrate that tripeptidyl peptidase II (TPP2) is essential for cerebrovascular homeostasis in adult female mice by orchestrating intracellular Ca²⁺ distribution and local estrogen biosynthesis. Specifically, TPP2 deficiency triggers a vicious cycle of Ca²⁺ imbalance and estrogen deficiency, thereby disrupting the anticipatory unfolded protein response (UPR). This disruption consequently drives aberrant pexophagic flux and ultimately depletes ether-linked phosphatidylcholine (PC-O)—the essential building block of endothelial cell (EC) membranes. In addition, we identified PC-O as a key facilitator of choline uptake through FLVCR2 and therefore its deficiency causes subsequent choline depletion as well as significantly decreased acetylcholine (ACh). Consistent with the fact that choline is a building block of PC and that ACh is a primary driver of vascular dilation, TPP2 depletion leads to significant cerebrovascular degeneration characterized by narrowed lumens, decreased EC number, and abnormal aggregation of ECs within the blood vessel. Additionally, AAV-mediated specific expression of Far1 in ECs to promote ether-PC biosynthesis significantly increases cerebrovascular volume and diameter in hippocampi of adult female TPP2 knockout (T2KO) mice. Importantly, ectopic expression of Far1 in ECs not only significantly ameliorates memory impairment in global and conditional TPP2-depleted female mice, but also significantly improves memory performance of naturally aged female mice. In total, our findings establish TPP2 as a key determinant of cerebrovascular homeostasis in adult female mice, positioning it as a novel therapeutic candidate for the treatment of VaD and AD.

## INTRODUCTION

The cerebrovascular endothelium, which forms the blood-brain barrier (BBB), is a highly specialized interface between the blood and the central nervous system (CNS). Dysfunction of endothelial cells (ECs) is a critical factor in the pathogenesis of a wide range of neurological and systemic diseases, such as neurodegenerative diseases^1, 2^, cerebral small vessel disease (cSVD)^3–5^, neuroinflammatory and autoimmune diseases^6, 7^, and acute ischemic and hemorrhagic stroke^8, 9^. These conditions are predominantly age-related neurological cognitive disorders, including vascular dementia (VaD)^10, 11^, Alzheimer’s disease (AD)^10, 12, 13^, and frontotemporal lobar degeneration (FTLD)^13–15^.

Age-related cognitive impairment poses a leading global public health challenge and a severe socioeconomic burden. While AD is the primary cause of clinically diagnosed dementia in Western countries, cognitive impairment of cerebrovascular origin is the second most common cause globally and may be the predominant one in East Asia^16–18^. A consistent observation across diverse ethno-regional groups is that women have a higher prevalence of dementia than men, with a gradient of variation evident across geographic and socioeconomic settings^19^. One study estimated that the lifetime risk of developing AD at age 45 is approximately 20% for women, compared to 10% for men—effectively double the risk. By age 65, the risks for both sexes increase slightly, though the disparity between women and men remains notable^16, 20^. Despite the clinical overlap and mixed pathology between VaD and AD, sex differences in VaD present a more complex picture than those in AD. While men are at a higher risk of developing VaD at a younger age, women appear to be especially vulnerable in later life and tend to experience a more severe progression of the disease^21, 22^. These observations make it imperative to adopt sex-sensitive approaches in the prevention, diagnosis, and management of both VaD and AD.

Estrogen deficiency is a key factor in the sexually dimorphic pathogenesis of both AD and VaD^21, 23, 24^. Accordingly, studies reported that estrogen counteracts Aβ/Tau pathology and supports synaptic function in AD^24–26^, while in VaD, it acts primarily by providing cerebrovascular benefits such as improved blood flow, anti-inflammatory effects, and mitigated insulin resistance^25, 27^. Considering that cerebrovascular impairment is a common key driver of both AD and VaD, as well as a spectrum of other neurological disorders^10–13, 28^, estrogen deficiency may act as a shared pathogenic nexus, operating through induction of cerebrovascular damage in addition to other parallel or hierarchical mechanisms to contribute to both diseases^29^.

Estrogen production involves a dynamic, hypothalamic-pituitary-gonadal (HPG) axis-mediated interplay between systemic and local sources^30^. While the gonads supply cyclical systemic estrogen, the brain synthesizes it locally *de novo* via CYP19A1 in neurons and glial cells in regions like the hippocampus and cortex^31–36^. Notably, CYP19A1 is also expressed in cerebrovascular ECs, providing a direct mechanism for the local synthesis of 17β Estradiol (E2) at the neurovascular interface^37–39^. This local production is poised to play a critical role in cerebrovascular health and neuroprotection^35, 40, 41^. Thus, impaired local estrogen synthesis may underlie the cerebrovascular impairment following the menopausal decline in systemic levels^24, 42, 43^. This mechanism may explain the heightened susceptibility of the female brain to both VaD and AD with aging^25, 44–47^.

Estrogen has long been recognized for its broad neuroprotective effects, which are relevant to both VaD and AD^43, 48^. This understanding is supported by large-scale observational studies, such as the Cache County Study, which consistently reported a more than 30% reduction in the risk of AD among women who initiated ERT/MHT around menopause^49–52^. This evidence fostered a powerful narrative of ERT as a preventive “shield” against cognitive decline. However, due to associated risks like breast cancer, stroke, and blood clotting, ERT is currently not approved for use in preventing or treating VaD and AD, its clinical use is now solely confined to the management of menopausal symptoms^53, 54^. In view of this, the future of ERT is advancing toward greater precision and enhanced safety. A key strategy involves developing tissue-selective agents, such as selective estrogen receptor modulators (SERMs), which are engineered to mimic estrogen’s benefits in tissues like bone and brain while blocking its action in the breast and uterus^55–59^. A parallel approach aims to boost or restore local neurosteroidogenesis, thereby maximizing protective effects within the brain while minimizing peripheral risks^60–63^. Both pathways seek to deliver therapeutic benefits without the associated risks of traditional ERT.

In this study, we investigated the role of TPP2 in maintaining estrogen homeostasis in cerebrovascular ECs and perivascular astrocytes of adult female mice. We found that both global and selective depletion of TPP2 in either cerebrovascular ECs or perivascular astrocytes result in severe local estrogen depletion and marked cerebrovascular degeneration compared with controls. Mechanistically, we found that TPP2 depletion disrupts the anticipatory UPR signaling pathway in cerebrovascular ECs. Specifically, it triggers a vicious cycle between Ca²⁺ imbalance and estrogen deficit, which consequently drives aberrant pexophagy and ultimately results in PC-O deficiency. Crucially, PC-O deficit causes choline uptake impairment and ACh deficit and ultimately leads to cerebrovascular degeneration and memory loss of female mice. These findings were confirmed by rescue experiments demonstrating that AAV-mediated ectopic expression of either CYP19A1 or Far1 is able to significantly increase hippocampal vascular volumes and diameters of TPP2-depleted and naturally aged female mice. In addition to that, ectopic expression of Far1 significantly restores learning and memory performance in TPP2-depleted and naturally aged female mice. In conclusion, this study reveals a novel mechanism underlying cerebrovascular degeneration in adult female mice and provides a more precise and safer therapeutic strategy for neurodegenerative diseases like VaD and AD.

## RESULTS

### Cerebrovascular degeneration is significantly accelerated in T2KO adult female mice compared with control mice

TPP2 has been demonstrated to function in the cytosol through both enzymatic and non-enzymatic mechanisms^64, 65^. Importantly, it has been implicated in human neurological diseases^64^. However, the underlying molecular mechanisms remain incompletely understood. To elucidate the underlying mechanisms of TPP2 in these diseases, we generated T2KO mice. Immunofluorescence staining of mouse brain slices showed that the precapillary arteries in 12-month-old female T2KO mice display abnormal morphology, mainly characterized by narrowed lumens, along with a decrease in EC number and their abnormal agglomeration within the vessels (Fig. 1A). In contrast, no significant morphological alterations were observed in 3-month-old female T2KO mice, or in male T2KO mice at either 3 or 12 months of age (Fig.1 A). Using bright field Micro-Optical Sectioning Tomography (MOST) imaging technique, we observed a marked reduction in vascular density in integrated holographic images from transverse and coronal sections, encompassing cortical and hippocampal areas (Fig. 1B). Statistical analysis of high-magnification images confirmed that cerebral capillaries in 12-month-old female T2KO mice exhibit a marked reduction in volume and diameter within both the cortex and hippocampus, whereas the length density and segment length show no significant change (Fig.1 B3). Consistent with immunofluorescence staining and MOST imaging, Masson, Elastica van Gieson (EVG), and Alkaline phosphatase (ALP) staining also showed that pathological alterations in the precapillary arteries of 12-month-old female T2KO mice, including luminal narrowing, reduction and agglomeration of ECs, and aberrant expression and distribution of collagen fibers and elastic fibers within the vessel wall (Fig. 1C). Furthermore, ultrastructural analysis by transmission electron microscopy (TEM) identified abnormal precapillary arteriole morphology in 12-month-old female T2KO mice, featuring luminal narrowing, vessel deformation, and astrocyte hypertrophy, thereby validating the multi-modal imaging data (Fig. 1D). To identify which type of vascular cells play a primary role in this pathological process, we prepared cell specific conditional T2KO mice. Interestingly, conditional knockout of TPP2 gene in either vascular ECs (Cdh5-Cre/TPP2^fl/fl^) or perivascular astrocytes (GFAP-Cre/TPP2^fl/fl^) induces cerebrovascular degeneration in 12-month-old female mice. In contrast, knockout of TPP2 in neurons (CamK2-Cre/TPP2^fl/fl^), microglia (TREM119-Cre/TPP2^fl/fl^), or neural stem cells (Nestin-Cre/TPP2^fl/fl^) does not produce this sex- and age-dependent degenerative effect (Fig. 1E). To determine the alteration of cell type composition of the mouse hippocampus in the absence of TPP2, we performed single-cell RNA sequencing (scRNA-seq) analysis. Our data revealed a marked reduction in the EC population in 12-month-old female mice following global TPP2 depletion (Fig. 1F-1G). Notably, this EC population is identified by a five-gene signature, including three established estrogen-regulated genes: CD34, Kdr, and Cldn5.

**Fig. 1.**
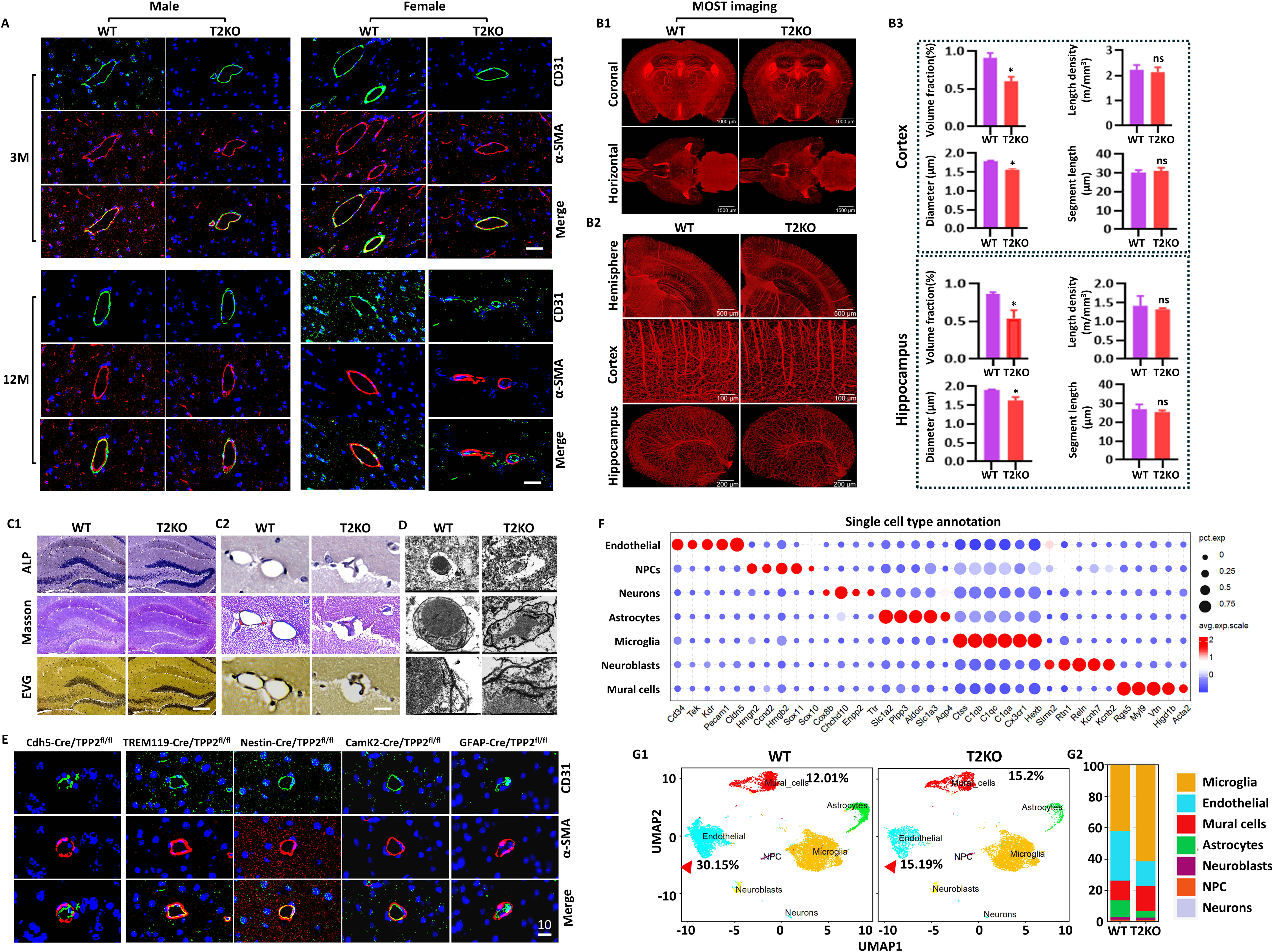
Cerebrovascular degeneration is accelerated in TPP2-depleted adult female mice compared with controls. **(A)** Immunofluorescence staining of mouse brain slices revealed that the precapillary arteries of 12-month-old female T2KO mice exhibit abnormal morphology, primarily characterized by narrowed lumens, as well as reduction and mislocalization of ECs within the vessels. In contrast, no significant morphological changes were observed in 3-month-old female T2KO mice, or in male T2KO mice at either 3 or 12 months of age. Representative results from one of three independent experiments are shown. scale bar: 10 µm. **(B)** Bright-field MOST (Micro-Optical sectioning tomography) imaging of mouse brain vascular atlas showed that the diameter and volume of cerebral capillaries in 12-month-old female T2KO mice are significantly reduced. **(B1)** Integrated holographic images of transverse and coronal sections and **(B2**) images of cortical and hippocampal regions revealed a visible marked reduction in vascular density. **(B3)** Statistical analysis confirmed that the volume and diameter of cortical and hippocampal capillaries in 12-month-old female T2KO mice are significantly reduced, while length density and segment length exhibit no significant change (n = 4, \**p* < 0.05, ns: not significant). **(C)** Masson, Elastica van Gieson (EVG) and Alkaline phosphatase (ALP) staining results are consistent with those from immunofluorescence staining and MOST. The precapillary arteries of 12-month-old female T2KO mice exhibit abnormal morphology, primarily characterized by narrowed lumens, reduction and mislocalization of ECs within the vessels (by ALP), as well as abnormal expression and distribution of collagen fibers (blue by Masson, red by EVG) and elastic fibers (black by EVG) on the blood vessel wall. Representative results from one of three independent experiments are shown. C1 scale bar: 200 µm, C2 scale bar: 10 µm. **(D)** Transmission electron microscopic (TEM) imaging showed that the precapillary arteries of 12-month-old female T2KO mice exhibit abnormal morphology, primarily characterized by narrowed lumens, as well as deformation of blood vessels and hypertrophy of astrocytes, corroborating the findings across multiple modalities. Zoom in from top to bottom sequentially. Representative results from one of three independent experiments are shown. **(E)** Conditional knockout of TPP2 in vascular ECs (Cdh5-Cre/TPP2^fl/fl^) or astrocytes (GFAP-Cre/TPP2^fl/fl^) induces cerebrovascular degeneration in 12-month-old female mice. In contrast, knockout of TPP2 in neurons (CamK2-Cre/TPP2^fl/fl^), microglia (TREM119-Cre/TPP2^fl/fl^), or neural stem cells (Nestin-Cre/TPP2^fl/fl^) does not produce this gender and age dependent degenerative effect. **(F)** Multi-group dotplot displaying the single cell type annotation in scRNA-seq analysis of mouse hippocampus. The horizontal axis represents the feature genes for identifying the cell types and the vertical axis lists the identified cell types. Circle size and colour indicate percent expression (Pct. Exp.) and expression scale (Exp. Scale) of the marker genes, respectively. **(G)** The identification and proportional composition of cell types in scRNA-seq analysis of mouse hippocampus are showcased through two visualizations: a Uniform Manifold Approximation and Projection (UMAP) graph (**G1**), which depicts cell clustering, and a stacked bar chart (**G2**), which quantifies the percentage abundance of each type. Our data indicate that the percentage of ECs identified by a five-gene signature decreases by nearly 50% when TPP2 gene is missing.

### Cerebrovascular degeneration in T2KO female mice is closely associated with choline uptake impairment

To identify the molecular mechanisms underlying TPP2 depletion-induced cerebrovascular degeneration, we established a T2KO bEnd.3 cell line and conducted a global metabolomic analysis. This approach identified a severe shortage of choline and its derivatives as a key metabolic consequence of TPP2 depletion (Fig. 2A). In brief, TPP2 depletion induces a distinct metabolic separation between wild-type (WT) and T2KO bEnd.3 cells, as shown by an Orthogonal Partial Least Squares Discriminant Analysis (OPLS-DA) model with excellent fit and predictive power (R2Y = 0.999, Q2 = 0.819) and with no overfitting (Fig. 2A1). Notably, PC and choline rank as the top two significantly deficient metabolites driving this separation in the Variable Importance in Projection (VIP) score plot (Fig. 2A2). We then conducted an ACh quantification assay in both bEnd.3 cells and primary mouse brain-derived vascular cells. Consistent with data of the global metabolomic analysis, TPP2 depletion resulted in a significant reduction in ACh levels in both models (Fig. 2B).

**Fig. 2.**
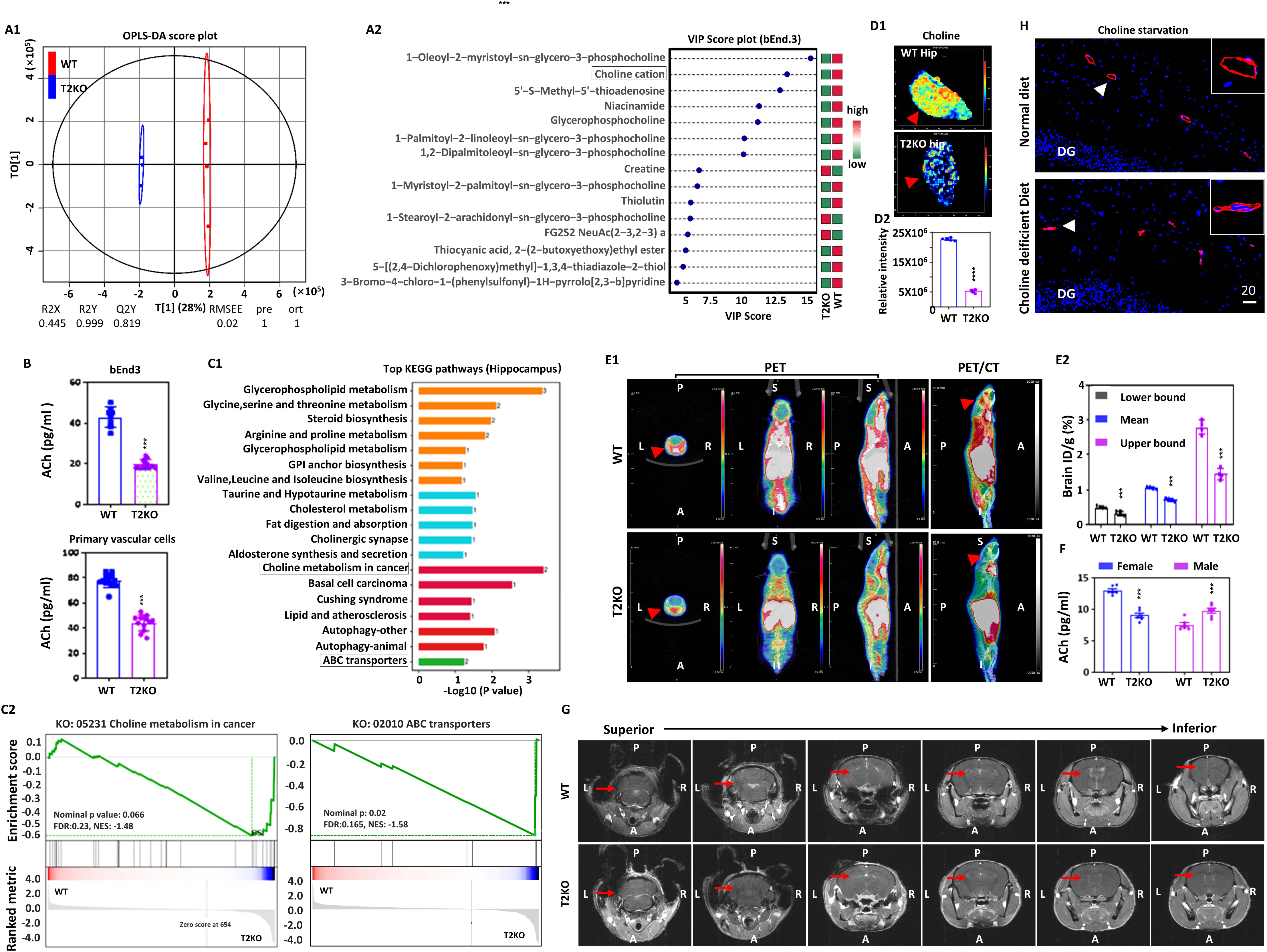
Cerebrovascular degeneration in TPP2-depleted female mice is closely associated with choline uptake impairment. **(A)** Global metabolomic analysis of bEnd.3 cells revealed that TPP2 depletion leads to a shortage of choline and its derivatives. **(A1)** OPLS-DA score plot illustrating the metabolic separation between WT and T2KO bEnd.3 cells. The horizontal axis corresponds to the first predictive component (t[1]), while the vertical axis represents the first orthogonal component (to[1]). The value indicated on the t[1] axis shows that this component accounts for 28% of the total variance in the original dataset. Colored dots represent samples from each group, and the ellipse denotes the 95% confidence interval. R2X (Explanation rate of X): 0.445; R2Y (Explanation rate of Y): 0.999 (> 0.6: good fitting); Q2 (Predictive ability of the model): 0.819 (> 0.5: strong predictive ability); R2Y-Q2 = 0.18 (< 0.3: No overfitting); RMSEE (Root Mean Squared Error of Estimation): 0.02 (< 0.1); pre (Number of predictive components extracted by the model): 1; ort (Number of orthogonal components extracted by the model): 1. **(A2)** VIP score plot illustrating the top metabolites driving the separation of metabolic profiles in A1. The x-axis corresponds to the VIP scores of the metabolites, while the y-axis displays the metabolite names ranked in descending order of VIP score from top to bottom. Metabolites with increased mean abundance are highlighted in red, whereas those with decreased mean abundance are indicated in blue. The significant reduction in choline is emphasized by a square enclosing its name to facilitate identification. **(B)** The ACh (Ach) quantification assay conducted in both bEnd.3 cells and primary mouse vascular cells demonstrated that TPP2 depletion results in a significant reduction in Ach levels (bEnd.3: n = 8, \*\*\**p* < 0.001; primary vascular cells: n = 12, \*\*\**p* < 0.001). **(C)** Spatial metabolomic analysis of mouse hippocampus revealed that TPP2 depletion leads to altered activity of choline metabolic and transport pathways. **(C1)** KEGG pathway enrichment analysis highlighting the most significantly impacted biological processes (BPs) following TPP2 depletion. The statistical significance (-log10(p-value)) of the enriched BPs is plotted on the x-axis, with the corresponding BP names displayed on the y-axis; the detected KEGG pathways are annotated by color to denote pathway type and by column size to represent the number of differentially expressed genes (DEGs). The choline metabolism and transport pathways are denoted by a square around their names for easy identification. **(C2)** GSEA revealed enriched gene sets for choline metabolism and ATP-binding cassette (ABC) transporters in a spatial metabolomics dataset of the mouse hippocampus, indicating altered activity in these pathways. **(D)** Spatial metabolomic analysis revealed that TPP2 depletion leads to a shortage of choline in female mouse hippocampus**. (D1)** Schematics illustrating spatial distribution of choline in the healthy control (WT) and T2KO samples. **(D2)** Statistical analysis showing that choline is very significantly decreased in hippocampi of T2KO mice (n = 6, \*\*\**p* < 0.001). **(E)** PET/CT imaging revealed a marked decrease in choline uptake within the brains of female mice following TPP2 depletion. **(E1)** PET/CT images of the transverse sections of the brain along with the coronal and sagittal sections of a mouse body at 60 minutes after injection of N-[^11^C]methyl choline chloride, illustrating choline uptake dynamics in the brain are severely impaired in the absence of TPP2 function. Representative images from one of three independent experiments are displayed. Red arrow highlighting the brain area; P: posterior; A: anterior; S: Superior; I: Inferior. **(E2)** Statistical analysis showing that choline uptake is very significantly decreased in brains of female T2KO mice (n = 4, \*\*\**p* < 0.001). % ID/g: Radioactive count per gram of tissue/total radioactive count injected X100%; Upper bound: The maximum intensity value within the image; Lower bound: The minimum intensity value within the image; Mean: The average intensity value across the entire image**. (F)** Analysis of cerebral Ach levels demonstrated that TPP2 depletion leads to a significant reduction in Ach in female mice, but not in males, indicating a sex-specific effect. **(G)** NMR/MRI dynamic imaging following injection of gadoterate meglumine salt solution showed no clear BBB permeability-related vasogenic edema in TPP2 depleted mice compared to WT controls. (**H)** Feeding a choline-deficient diet (CDD) to one-month-old female WT mice for one month recapitulates the cerebrovascular degeneration observed in the hippocampal region of female T2KO mice. This degeneration is characterized by narrowed vascular lumens, reduced EC numbers, and abnormal EC localization within vessels. A representative image from one of the six mice is shown, with an inset providing a magnified view of the area indicated by the arrow.

To determine whether choline is deficient in the hippocampi of T2KO mice, we further conducted unbiased spatial metabolomics analysis. Our data identified a global reduction in choline throughout the hippocampus (Fig.2 D1). Statistical analysis showed that choline is very significantly decreased in hippocampi of female T2KO mice (Fig. 2D2). Both Kyoto Encyclopedia of Genes and Genomes (KEGG) and Gene Set Enrichment Analysis (GSEA) enrichment analyses of the acquired data demonstrate significant alterations in choline metabolism and ATP-binding cassette (ABC) transporters (Fig. 2C), validating that TPP2 depletion disrupts these critical metabolic and transport processes.

To investigate the in vivo choline availability and its transport kinetics, we performed Positron Emission Tomography/Computed Tomography (PET/CT) imaging analysis. PET/CT images of the transverse sections of the brain along with the coronal and sagittal sections of a mouse body at 60 minutes after injection of N-[^11^C]methyl choline chloride showed that choline uptake dynamics in the brain are severely impaired in the absence of TPP2. Compared with WT controls, the % ID/g value of radioactive choline in female T2KO mice is reduced by nearly half (Fig. 2E). To determine whether impaired choline uptake leads to altered cerebral ACh level in T2KO mice, we measured ACh concentrations in whole-brain homogenates from both male and female mice. Our data demonstrated that TPP2 depletion causes a significant decrease in cerebral ACh in females, but an increase in males, indicating a sex-specific effect (Fig. 2F).

To further determine whether the choline deficiency in female T2KO mice is caused by changes in BBB, we conducted Nuclear Magnetic Resonance (NMR/MRI) imaging on the BBB of T2KO mice. Although TPP2 depletion resulted in an ∼50% loss of the identified EC population, the NMR/MRI dynamic imaging following injection of gadoterate meglumine salt solution showed no accompanying increase in BBB permeability or vasogenic edema in T2KO mice (Fig. 2G). To assess whether choline deficiency accelerates cerebrovascular degeneration, female mice were maintained on a choline-deficient dietary (CDD) regimen. Interestingly, long-term feeding of CDD recapitulates the cerebrovascular degeneration observed in the hippocampal region of female T2KO mice (Fig. 2H).

### Choline uptake impairment in TPP2-depleted cerebrovascular ECs is associated with the Ca²⁺-E2-PC-O signaling pathway

To investigate the molecular mechanism of impaired choline uptake, we performed scRNA-seq on hippocampal tissues from female T2KO mice and their WT counterparts. KEGG pathway enrichment analysis of the scRNA-seq data identified the top 20 most significantly altered biological processes (BPs) in ECs following TPP2 depletion (Fig. 3A1). We further performed Gene Set Variation Analysis (GSVA) to derive hallmark BPs, followed by a correlation analysis to investigate their interrelationships (Fig. 3A2). This integrated approach revealed significant positive correlations between choline metabolism and several signaling pathways—specifically estrogen, Ca²⁺, ether lipid, and glycerophospholipid metabolism (Fig. 3A). The strength of these linear relationships, quantified by Pearson correlation coefficient (PCC) values (where +1 and -1 represent perfect positive and negative correlations, respectively), range from 0.5 to 0.8, suggesting a potential synergistic role among these pathways in choline regulation (Fig. 3A).

**Fig. 3.**
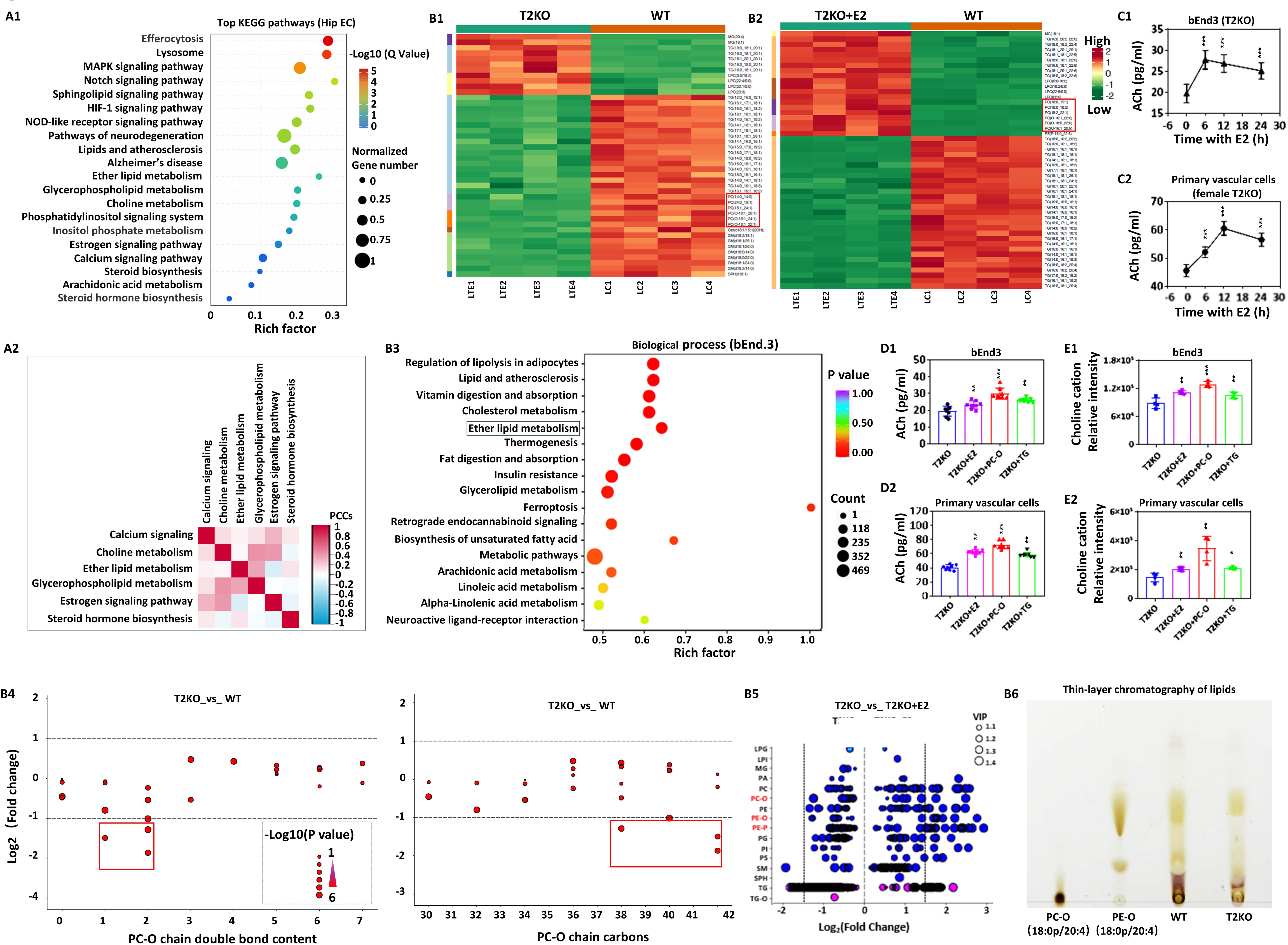
Choline uptake impairment in TPP2-depleted cerebrovascular ECs is associated with the Ca²⁺-E2-PC-O signaling pathway. **(A)** scRNA-seq analysis of hippocampal ECs revealed that choline dysregulation is closely associated with the Ca²⁺, E2, and PC-O signaling pathways. **(A1)** KEGG pathway enrichment analysis, performed on scRNA-seq data, highlights the most significantly affected BPs in hippocampal ECs after TPP2 depletion. The rich factor of each enriched pathway is shown on the x-axis, while the corresponding pathway names are listed on the y-axis. Significantly enriched KEGG pathways are color-coded according to their statistical significance (–log₁₀(Q value)), and the circle size represents the number of normalized DEGs. The ether lipid metabolism pathway is highlighted with a square around its name for easy identification (Rich factor > 0.25, *p* < 0.01)**. (A2)** Correlation analysis of hallmark BPs derived from GSVA in panel (A1) reveals significant positive associations between choline metabolism and multiple signaling pathways-including estrogen signaling, Ca²⁺ signaling, ether lipid metabolism, and glycerophospholipid metabolism-suggesting a potential synergistic role in the regulation of choline levels. The strength of these linear relationships was quantified using the PCC (Pearson Correlation Coefficient) value, where a value of +1 denotes a perfect positive correlation and –1 indicates a perfect negative correlation. **(B)** Lipidomic analysis of bEnd.3 cells revealed that TPP2 depletion causes a deficiency in PC and PC-O species, which could be reversed by E2 treatment. **(B1)** Clustered heatmap displaying the overall distribution and relative abundance of differential lipid species in WT and T2KO samples. The significant reduction in PC and PC-O at absence of TPP2 is emphasized by a square enclosing their names to facilitate identification (n = 4, VIP > 1, *p* < 0.001). **(B2)** Clustered heatmap displaying the overall distribution and relative abundance of differential lipid species in WT and T2KO samples with E2 treatment for 6 hours. The results demonstrate that E2 rescues the deficiency in PC and PC-O lipids in T2KO bEnd.3 cells. The significant upregulation in PC and PC-O by E2 treatment is emphasized by a square enclosing their names to facilitate identification. For (B1) and (B2), the horizontal axis indicates the sample labels. The left vertical axis colour bar corresponds to lipid categories, and the right axis shows specific lipid names. Each colored cell represents the expression level of a lipid, with darker shades corresponding to higher values. Red indicates increased abundance, while green denotes decreased levels of differential lipids (n = 4, VIP > 1, *p* < 0.001). **(B3)** KEGG enrichment analysis of BPs, performed on lipidomic data, highlighting the most significantly affected BPs in bEnd.3 cells after TPP2 depletion. The rich factor of each BP is shown on the x-axis, while the corresponding BP names are listed on the y-axis. Significantly enriched BPs are color-coded according to their statistical significance *(p* value), and the circle size represents the number of DEGs. The ether lipid metabolism pathway is highlighted with a square around its name for easy identification (n = 4, Rich factor > 0.65, *p* < 0.01)**. (B4)** Scatter plots revealed a significant reduction in PC-O species containing fewer than three double bonds and long carbon chains (>36 carbons) within T2KO bEnd.3 cells. The x-axis represents the number of double bonds (left) or carbon atoms (right), while the y-axis displays the log₂(Fold change) values for the corresponding lipid classes. PC-O species significantly downregulated due to TPP2 depletion are highlighted in the red square (n = 4, |log₂(Fold Change)| > 1, *p* < 0.001). **(B5)** The scatter plot demonstrating that E2 significantly upregulates glycerophospholipids-specifically PC-O, ether-linked phosphatidylethanolamine (PE-O), and phosphatidylethanolamine plasmalogen (PE-P)-in T2KO bEnd.3 cells compared to control cells without E2 treatment (n = 4; |log₂(Fold Change)| > 1.5, VIP > 1, *p* < 0.001), while TG (triglyceride) levels remain largely unaffected by E2 supplementation, consistent with the findings shown in panels B1 and B2. The x-axis corresponds to log₂(Fold change), and the y-axis indicates the corresponding lipid classes. Ether-linked lipids are highlighted in red on the y-axis. **(B6)** Thin-layer chromatographic analysis of phospholipids indicates that TPP2 depletion leads to a significant reduction in PC-O, using 18:0p/20:4 PC and 18:0p/20:4 PE as reference standards. (**C)** Dynamic analysis of ACh (ACh) levels following E2 treatment in bEnd.3 cells and primary vascular cells of female mouse brains revealed that ACh upregulation peaks at 6 hours in bEnd.3 cells and at 12 hours in primary vascular cells after E2 treatment, respectively (n = 6, \*\*\**p* < 0.001). **(D)** Quantitative colorimetric analysis revealed that E2, PC-O, or TG significantly elevate ACh levels in both T2KO bEnd.3 cells and the vascular cells of female T2KO mice (n = 8, \*\*\**p* < 0.001, \*\**p* < 0.01). **(E)** Global metabolic analysis revealed that E2, PC-O, or TG significantly elevate choline levels in both T2KO bEnd.3 cells and the primary vascular cells of female T2KO mice (n = 4, \*\*\**p* < 0.001, \*\**p* < 0.01).

To validate these findings and further investigate the interplay among these signaling pathways in greater detail, we conducted lipidomics analysis on a T2KO bEnd.3 cell model with and without E2 treatment. Our data demonstrated that TPP2 depletion leads to a marked decrease in PC and PC-O species compared to WT controls, along with a subset of triacylglycerols (TGs) and sphingomyelins (SMs) (Fig. 3B). Interestingly, E2 treatment can significantly upregulate the levels of PC and PC-O species, whereas it does not markedly alter the levels of the affected TGs and SMs (Fig. 3B1-3B2). These findings indicate that E2 plays a critical role in maintaining the levels of PC and PC-O species in model ECs. Consistent with the findings in hippocampal ECs, KEGG enrichment analysis of the bEnd.3 cell model data likewise identified ether lipid metabolism as a significantly affected pathway following TPP2 depletion (Fig.3 B3). Furthermore, scatter plot and thin-layer chromatography (TLC) analysis confirmed that TPP2 depletion in bEnd.3 cells significantly reduces a specific subset of PC-O species, characterized by fewer than three double bonds and long carbon chains (>36 combined carbons of 2 chains) (Fig. 3B1, 3B4-3B6). It is noteworthy that E2 treatment rescues the reduced PC-O species in T2KO cells, yet these induced species were characterized by long carbon chains (>36 combined carbons) and ≥4 double bonds (Fig. 3B2 and 3B5), while the levels of TGs and SMs remain largely unaltered (Fig. 3B2 and 3B5).

Subsequently, we investigated the regulatory roles of Ca²⁺, E2, and PC-O in choline uptake. To address this objective, we first characterized the temporal dynamics of ACh levels after E2 administration in bEnd.3 cells and primary vascular cells. We observed that ACh peaks at 6 hours in bEnd.3 cells and at 12 hours in primary vascular cells after E2 treatment (Fig. 3C). We then further compared the effects of E2, PC-O, and TG on ACh levels. Quantitative colorimetric assays revealed that all treatments significantly increase ACh levels in both T2KO bEnd.3 cells and primary T2KO vascular cells from female mice, with PC-O showing the strongest effect (Fig. 3D). To confirm that ACh levels are closely linked to choline levels, we conducted a global metabolic analysis. The results showed that E2, PC-O, and TG significantly elevate choline levels in both T2KO bEnd.3 cells and primary T2KO vascular cells from female mice (Fig. 3E). Collectively, these data indicate that Ca²⁺, E2, and PC-O operate within the same signaling pathway for choline uptake regulation (Fig. 3C, 3D and 3E).

### The Ca²⁺ imbalance and E2 deficit form a vicious cycle via anticipatory UPR pathway in TPP2-depleted cerebrovascular ECs

Given the regulatory roles of Ca²⁺, E2, and PC-O on choline uptake, we first examined the integral relationship between Ca²⁺ and E2 in the absence of TPP2. Two-photon imaging of hippocampal slices loaded with Fluo-4 AM uncovered in situ reduced cytosolic Ca²⁺ levels in vascular cells of the CA1 and DG regions upon TPP2 depletion (Fig. 4A). Additional GSEA analysis performed on scRNA-seq data in hippocampal ECs revealed enriched gene sets for Ca²⁺ signaling pathway, indicating altered activity in this pathway upon TPP2 depletion (Fig. 4B). Moreover, genetically encoded Ca²⁺ indicators (GECIs)-based measurement revealed that TPP2 depletion in primary cerebrovascular ECs significantly decreases cytosolic Ca²⁺ while increasing endoplasmic reticulum Ca²⁺, and this phenotype can be rescued by E2 treatment (Fig. 4C). These results implicate a strong link between estrogen deficiency and dysregulated Ca²⁺ ion distribution (cytosol vs. ER) in T2KO cerebrovascular ECs.

**Fig. 4.**
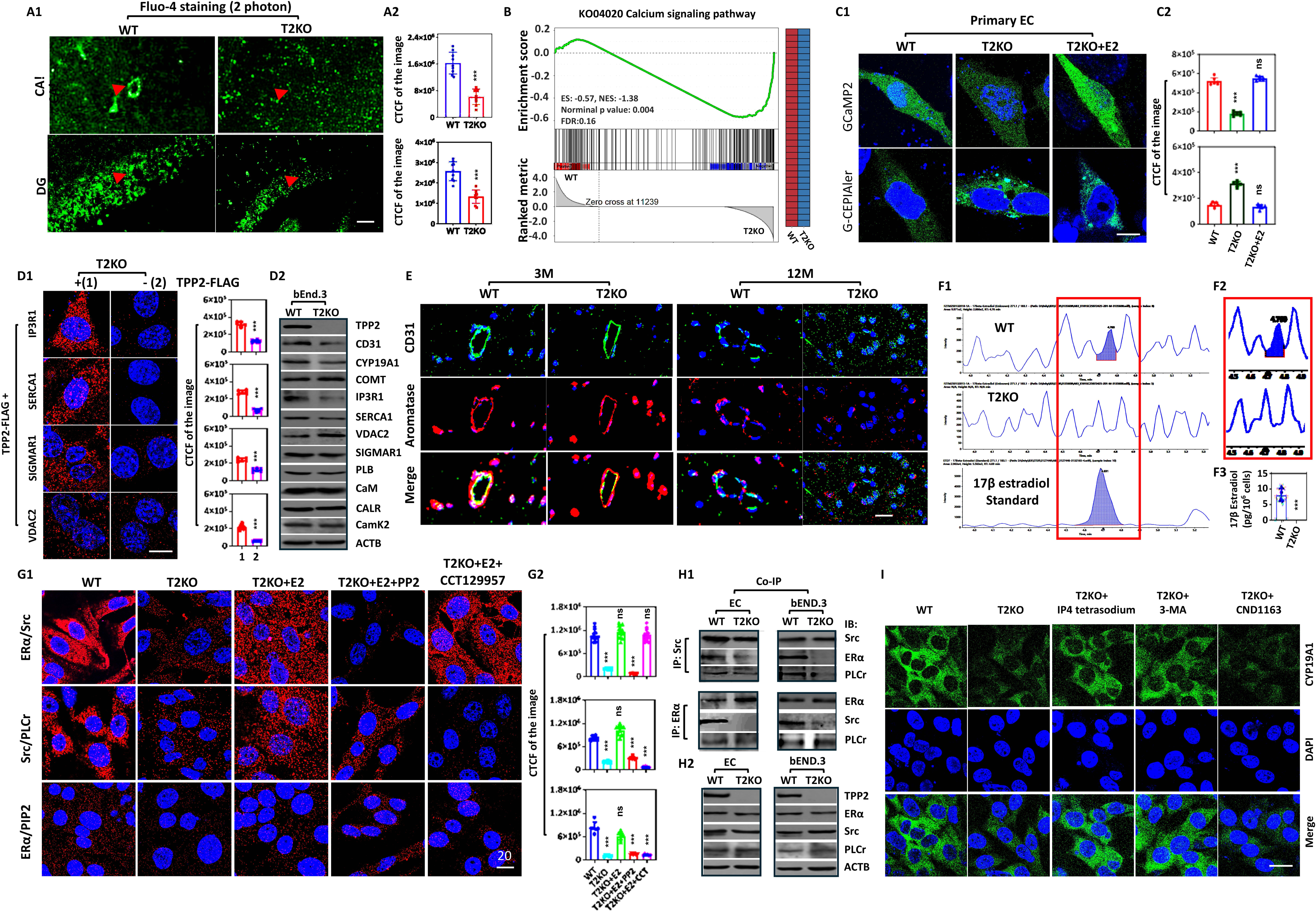
The Ca^2+^ imbalance and estrogen deficit form a vicious cycle via anticipatory UPR pathway in TPP2-depleted cerebrovascular ECs. **(A)** Two-photon imaging of hippocampal slices loaded with Fluo-4 AM revealed reduced cytosolic Ca²⁺ levels in vascular cells of the CA1 and DG regions at absence of TPP2. **(A1)** Representative images of Ca^2+^ in CA1 (upper) and DG (lower) regions. **(A2)** Statistical graphs of CTCF of the images. n = 4, ***p ≤ 0.001. Scale bar, 10 µm**. (B)** GSEA analysis performed on scRNA-seq data in hippocampal ECs at absence of TPP2 revealed enriched gene sets for Ca²⁺ signaling pathway, indicating altered activity in this pathway. **(C)** Genetically encoded Ca²⁺ indicators (GECIs)-based measurement revealed that TPP2 depletion in primary cerebrovascular ECs significantly decreases cytosolic Ca²⁺ while increasing endoplasmic reticulum Ca²⁺, and this phenotype can be rescued by E2 treatment. **(C1)** Representative images of cytosolic Ca^2+^ detected by GCaMP2 (upper) and ER Ca^2+^ detected by G-CEPIA1er (lower)**. (C2)** Statistical graphs of CTCF of the images. n = 6, ***p ≤ 0.001. Scale bar, 10 µm**. (D)** In bEnd.3 cells, TPP2 interacts with IP3R1, SERCA1, SIGMAR1, and VDAC2, and its depletion leads to the downregulation of IP3R1 and CYP19A1. **(D1)** Representative images of PLA and their statistical graphs of CTCF of the images. n = 6, ***p ≤ 0.001. Scale bar, 10 µm**. (D2**) Immunoblot analysis of the protein levels of IP3R1, SERCA1, SIGMAR1, VDAC2, and other related proteins. The results revealed a significant downregulation of IP3R1 and CYP19A1 upon TPP2 depletion. Data from one of three independent experiments are shown. **(E)** Immunofluorescence staining revealed significantly reduced levels of both CYP19A1 and CD31 in hippocampal vascular ECs of 12-month-old T2KO mice compared to WT controls. In contrast, no significant change was observed in 3-month-old mice. Shown is a representative result from three independent experiments. **(F)** Targeted metabolomic profiling of steroid hormones in primary vascular cells revealed that E2 was completely depleted in the absence of TPP2. Representative images of base peak **(F1)** and extracted ion chromatograms **(F2)** from four independent samples**. (F3)** Statistical graph of identified E2 showing that in WT vascular cells it is up to 10 pg per million cells, whereas it is undetectable in T2KO cells. n = 4, ***p ≤ 0.001. **(G)** PLA results indicated a severe disruption of the anticipatory UPR in TPP2-depleted primary ECs, which was restored by E2 treatment. This restoration was completely blocked by either Src inhibitor PP2 or PLCγ inhibitor CCT129957. Representative images of PLA **(G1)** and their statistical graphs of CTCF of the images **(G2)** are shown. n = 10, ***p ≤ 0.001. Scale bar, 20 µm. **(H)** Co-IP combined with immunoblot showed the interaction of the components of anticipatory UPR signaling pathway in both bEnd.3 cells and primary ECs is disrupted under TPP2 depletion. **(H1)** Immunoblot analysis of precipitates showed the decoupling among Src, ERα, and PLCγ under TPP2 depletion under TPP2 depletion. **(H2)** Immunoblot analysis of homogenates revealed no significant change in the protein levels of Src, ERα, and PLCγ of T2KO samples. Data from one of three independent experiments are shown. **(I)** In cerebrovascular ECs, TPP2 depletion leads to CYP19A1 deficiency via Ca²⁺ imbalance-mediated dysregulation of autophagic flux. This deficiency is exacerbated by CND1163, a SERCA activator that worsens cytosolic Ca²⁺ deficit. Conversely, it is rescued by treatment that elevates cytosolic Ca²⁺ (e.g., D-myo-inositol-1,2,4,5-tetrakisphosphate tetrasodium: IP4 tetrasodium, an IP3R1 activator) or treatment that inhibits autophagic flux (e.g., 3-methyladenine: 3-MA). Representive images from 12 independent experiments are shown. Scale bar, 20 µm.

To investigate how TPP2 depletion disrupts Ca²⁺ homeostatic distribution between the cytosol and ER and whether it concurrently induces estrogen deficiency in cerebrovascular ECs, we performed proximity ligation assay (PLA) and immunofluorescence staining. The PLA results demonstrated a direct physical interaction between TPP2 and IP3R1, SERCA1, SIGMAR1, and VDAC2 (Fig. 4D1). This suggests that loss of TPP2 may impair the function or stability of these Ca²⁺ regulatory proteins. Consistent with this possibility, TPP2 depletion led to a significant reduction not only in CYP19A1 but also in IP3R1 levels in bEnd.3 cells (Fig. 4D2). In consistence with this, immunofluorescence staining revealed a severe depletion of CYP19A1 in hippocampal vascular ECs of 12-month-old T2KO mice, whereas the CYP19A1 levels in 3-month-old mice remain largely unchanged upon TPP2 depletion (Fig. 4E). To corroborate our immunofluorescence findings, we performed targeted metabolomic profiling of steroid hormones in primary cerebrovascular cells (prepared from isolated cerebral vasculature) and immunoblot analysis of CYP19A1 in purified cerebral ECs and astrocytes from their corresponding conditional T2KO mice. Metabolomic analysis revealed that E2 levels were undetectable following either global or cell-specific TPP2 depletion, whereas WT controls maintained approximately 10 pg of estrogen per million cells (Fig. 4F). Consistent with this, immunoblotting showed a significant reduction in CYP19A1 levels in either purified cerebral ECs or astrocytes upon cell-specific depletion of TPP2 (Supplementary Fig. 3A).

It has been reported that estrogen regulates Ca²⁺ homeostasis through the anticipatory UPR signaling pathway^66, 67^. To test the integrity of this pathway, we performed PLA and found that this pathway is severely disrupted in T2KO ECs due to E2 depletion (Fig. 4G). This conclusion is supported by the finding that E2 replenishment fully restores the pathway activity, an effect completely blocked by co-treatment with either the Src inhibitor PP2 or the PLCγ inhibitor CCT129957 (Fig. 4G). Furthermore, co-immunoprecipitation (co-IP) coupled with immunoblot analysis demonstrated that TPP2 depletion disrupts the interactions between key components of the anticipatory UPR pathway in both bEnd.3 cells and primary vascular cells (Fig. 4H). Specifically, TPP2 deficiency led to the decoupling of Src, ERα, and PLCγ, despite causing no significant change in their total protein levels (Fig. 4H).

To further interrogate the mechanism underlying estrogen deficiency following TPP2 depletion, we specifically examined the link between cytosolic Ca²⁺ level and CYP19A1 expression in cerebrovascular ECs and astrocytes using immunostaining. Our results indicate that the CYP19A1 abundance is critically regulated by cytosolic Ca²⁺ levels (Fig. 4I). Supporting this notion, while the SERCA activator CND1163 exacerbated the CYP19A1 deficiency in T2KO ECs, the IP3R1 activator D-myo-inositol-1,2,4,5-tetraphosphate sodium (IP4 tetrasodium) prevented it (Fig. 4I). Intriguingly, inhibition of autophagy with 3-MA likewise prevented CYP19A1 deficit, implicating the Ca²⁺ imbalance-associated autophagic pathway in this process (Fig. 4I). Moreover, ectopic expression of catalytically inactive TPP2 mutants (TPP2 S449A and S449T) specifically rescued the CYP19A1 degradation in T2KO bEnd.3 cells (Supplementary Fig. 3B). Taken together, these data demonstrate that TPP2 maintains Ca²⁺ and estrogen homeostasis in an enzymatic activity independent manner, and its depletion triggers a vicious cycle between Ca²⁺ imbalance and estrogen deficiency within cerebrovascular ECs, thus potentially explaining the observed cerebrovascular degeneration in the absence of TPP2.

### The Ca²⁺-estrogen vicious cycle under TPP2 depletion leads to severe peroxisome deficit via an aberrant pexophagy flux and explains the decreased ether phospholipid production as shown above

Given that Ca²⁺, estrogen, and PC-O function within the same signaling pathway for choline uptake, with estrogen being critical for maintaining PC-O species levels in ECs (Fig. 3), we next investigated how the vicious cycle between Ca²⁺ imbalance and estrogen deficiency leads to a subsequent PC-O deficit. Considering the importance of peroxisome in PC-O biosynthesis, we performed two sets of experiments: one to assess peroxisomal abundance following TPP2 depletion, and the other to evaluate the effect of E2 and TG treatment on peroxisomal abundance. Our results revealed that TPP2 depletion causes a severe reduction in peroxisome abundance in both bEnd.3 cells and primary ECs, which is effectively prevented by treatment with either E2 or TG (Fig. 5A). To ensure robustness of these findings, we employed two different peroxisomal markers (pmScarlet-1_peroxisome_C1: PTS-Scarlet; pRRLSIN.cPPT.PGK-GFP-PTS1.WPRE: PTS-GFP), both of which yielded highly consistent results (Fig. 5A).

**Fig. 5.**
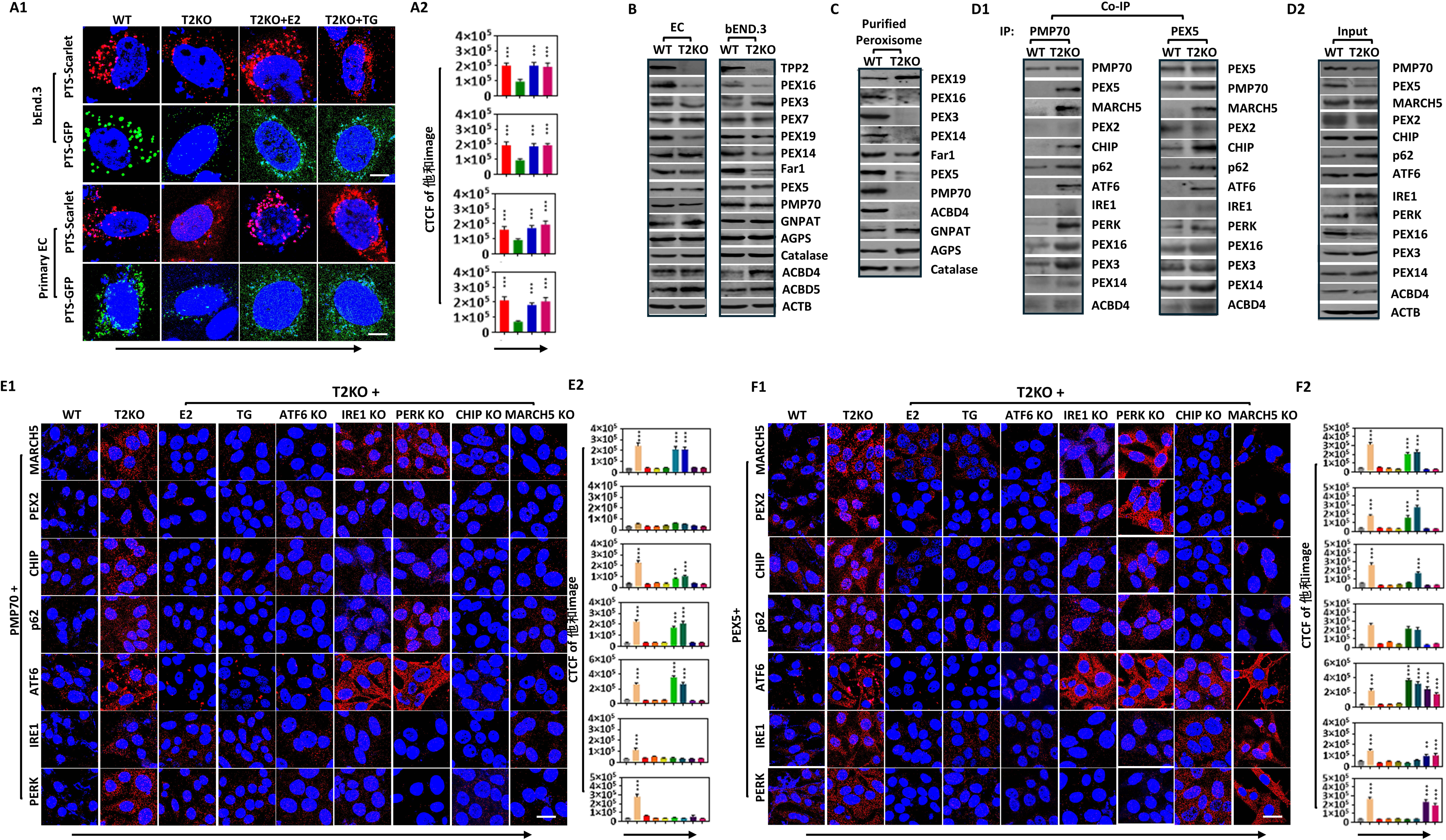
The Ca^2+^-estrogen vicious cycle under TPP2 depletion leads to severe peroxisome deficit via an aberrant pexophagy flux and explains the decreased ether phospholipid production as shown above. **(A)** TPP2 depletion caused a severe reduction in peroxisome abundance in both bEnd.3 cells and primary ECs, which was effectively prevented by treatment with either E2 or TG. **(A1)** Representative images of peroxisomes labeled by PTS-Scarlet (Red) and PTS-GFP (Green)**. (A2)** Statistical graphs of CTCF of the images. n = 12, ***p ≤ 0.001. Scale bar, 5 µm. **(B)**Immunoblot analysis of proteins involved in peroxisomal biogenesis, cargo import, membrane integrity, and matrix enzymes revealed that the levels of PEX16, Far1, PEX5, PMP70, and PEX19 are decreased in both bEnd.3 cells and primary vascular cells upon TPP2 depletion. No consistent differences were observed for the other proteins shown. **(C)** Analysis of isolated peroxisomes revealed that proteins for membrane assembly (e.g. PEX16) and cargo import (e.g. PMP70 and PEX5) are decreased or missing, whereas the levels of matrix enzymes (e.g. GNPAT, AGPS, and catalase) are surprisingly unaltered. **(D)** Co-IP combined with immunoblot identified the active pexophagy flux in T2KO bEnd.3 cells. **(D1)** Immunoblot analysis of precipitates determined the pexophagic complex components, which include PMP70/PEX5 (autophagic adaptor), MARCH5/CHIP (ubiquitin E3 ligase), p62 (autophagic receptor), ATF6/PERK (ER stress sensor), and PEX16/PEX3/PEX14/ACBD3. **(D2)** Immunoblot analysis of homogenates determined the overall expressin levels of the target proteins. **(E)** PLA data disclosed that the in situ interactions between PMP70 and other pexophagic complex components in T2KO bEnd.3 cells are strongly dependent on ATF6, CHIP, and MARCH5. Representative images of PLA **(E1)** and their statistical graphs of CTCF of the images **(E2**) are shown. n = 10, ***p ≤ 0.001, **p ≤ 0.01. Scale bar, 10 µm. **(F)** PLA revealed that the in situ interactions between PEX5 and other pexophagic complex components in T2KO bEnd.3 cells are strongly dependent on ATF6, but selectively dependent on CHIP and MARCH5. Representative images of PLA **(F1)** and their statistical graphs of CTCF of the images **(F2**) are shown. n = 10, ***p ≤ 0.001, **p ≤ 0.01. Scale bar, 10 µm. Treatment with either E2 or TG abolishes the pexophagic complex formation.

To investigate the mechanism underlying peroxisomal deficit in cerebrovascular ECs upon TPP2 depletion, we examined the expression of key proteins involved in peroxisomal biogenesis, cargo import, membrane integrity, and matrix enzyme function by immunoblotting. Our analysis revealed that TPP2 depletion consistently decreases the levels of PEX16, Far1, PEX5, PMP70, and PEX19 in both bEnd.3 cells and primary cerebrovascular cells, whereas no consistent differences for the other proteins were observed (Fig. 5B). We then isolated peroxisomes and endoplasmic reticulum and analyzed their protein content. In isolated peroxisomes, there is a specific loss or reduction of proteins critical for peroxisomal membrane assembly (e.g., PEX16) and cargo import (e.g., PMP70 and PEX5) under TPP2-deficient conditions, whereas the levels of matrix enzymes (e.g. GNPAT, AGPS, and catalase) are surprisingly unaltered (Fig. 5C). In contrast, the levels of PEX16, Far1, PEX5, PMP70, and PEX19 are elevated in the endoplasmic reticulum upon TPP2 depletion (data not shown). This suggests that these peroxisomal proteins accumulate in the endoplasmic reticulum and are not efficiently incorporated into newly formed peroxisomes derived from the ER. Further co-IP combined with immunoblot identified the active pexophagy flux in T2KO bEnd.3 cells (Fig. 5D). Specifically, immunoblot analysis of the precipitates identified key components of the pexophagic complex associated with the well-known autophagic adaptor PMP70/PEX5, such as ubiquitin E3 ligases MARCH5/CHIP, autophagic receptor p62, ER stress sensors ATF6/PERK, and peroxisomal membrane proteins PEX16/PEX3/PEX14/ACBD3 (Fig. 5D). To dissect the pexophagic flux in more details, we performed PLA to determine the in situ interactions among the pexophagic components. Our results confirmed that both pexophagic adaptors (PMP70 and PEX5) are involved in the pexophagic flux (Fig. 5E and 5F). Furthermore, the identified PMP70-mediated pexophagic flux is strongly dependent on ATF6, CHIP, and MARCH5 (Fig. 5E), whereas PEX5-mediated one is strongly dependent on ATF6, but selectively dependent on CHIP and MARCH5 (Fig. 5F). Most importantly, treatment with either E2 or TG abolishes the pexophagic complex formation (Fig. 5E).

### Ether phospholipid impacts choline incorporation via allosteric binding to FLVCR2

As abovementioned, Ca²⁺, estrogen, and PC-O function within the same signaling pathway for choline uptake, and the vicious cycle between Ca²⁺ imbalance and estrogen deficiency leads to a subsequent PC-O deficit, we next investigated the molecular mechanism underlying the regulation of choline uptake by PC-O. To this end, we performed an initial screening of multiple fluorescent choline analogs in order to select the most suitable probe for subsequent choline uptake measurements. We evaluated three specific fluorescent choline analogs, namely Choline-O-Cy2 (Cy2 fluorophore conjugated to the hydroxyl group), Choline-CO-Cy2 (Cy2 fluorophore attached to the carbon atom adjacent to the hydroxyl group), and Choline-CH3-NBD (NBD fluorophore linked to the methyl group). Our results revealed that the three probes exhibit a spectrum of uptake efficiency, ranging from the highest incorporation for choline-O-Cy2, to partial uptake for choline-CO-Cy2, and finally to a complete absence of uptake for choline-CH3-NBD in WT bEnd.3 cells (Fig. 6A). Crucially, the lack of detectable uptake for all three analogs in the absence of TPP2 (Fig. 6A) confirms that our assay measures specific, TPP2-dependent choline incorporation rather than background incorporation.

**Fig. 6.**
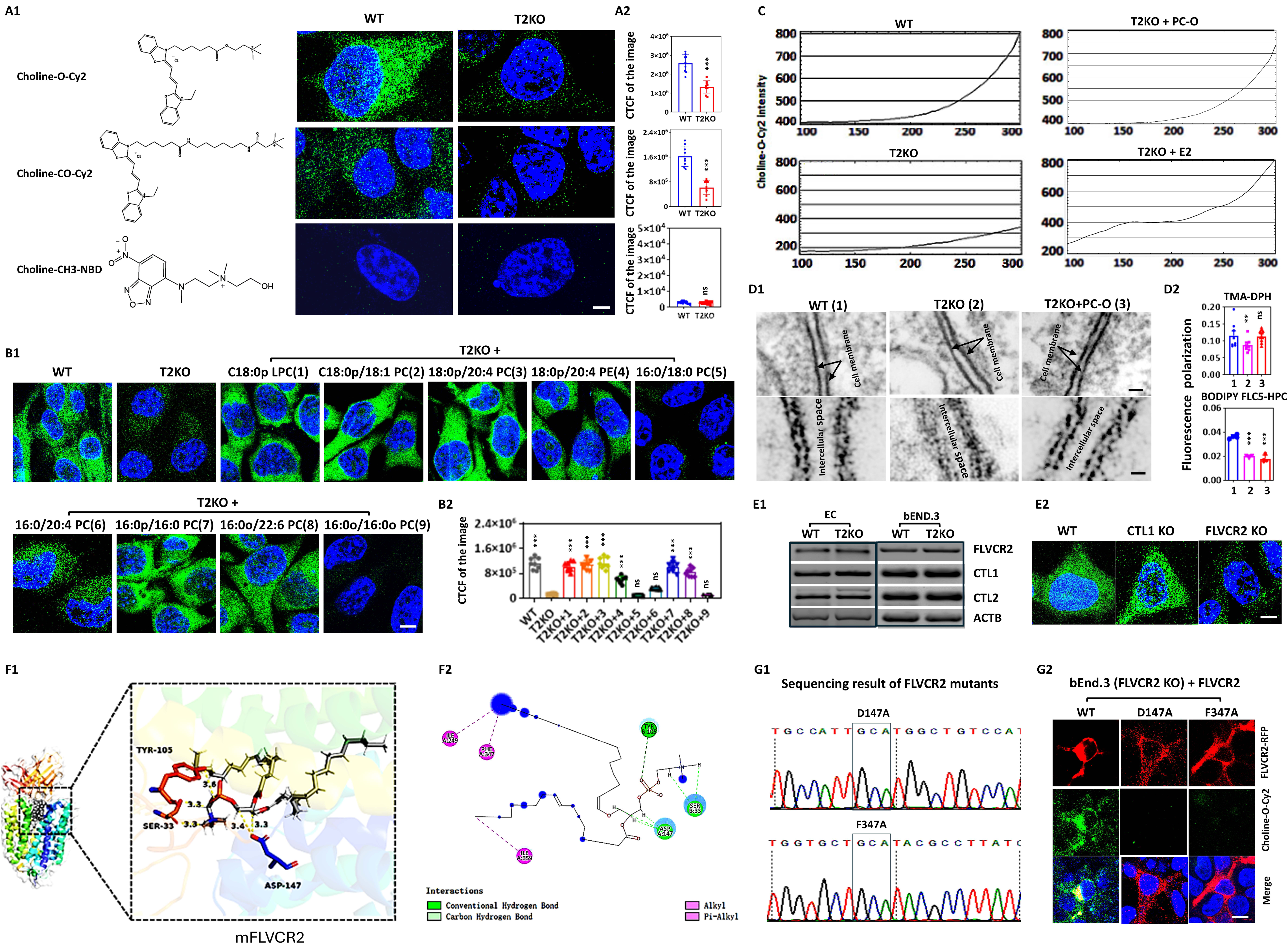
Ether phospholipids impact choline incorporation via allosteric binding to FLVCR2. **(A)** Screening of various fluorescent choline analogs via a choline uptake assay revealed that the version with a Cy2 fluorophore conjugated to the hydroxyl group (Choline-O-Cy2) exhibits the highest uptake efficiency. Conversely, NBD fluorophore linked to the methyl group (Choline-CH3-NBD) completely abolishes choline incorporation, and Cy2 fluorophore attached to the carbon atom adjacent to the hydroxyl group (Choline-CO-Cy2) results in partial inhibition. **(A1)** Illustration of the molecular structures of fluorescently labeled cholines **(left)** and representative images of incorporated cholines **(right). (A2)** Statistical graphs of CTCF of the images. n = 6, ***p ≤ 0.001. Scale bar, 2.5 µm. T2KO cells were used as a control to ensure that measured choline uptake was TPP2-dependent and to rule out nonspecific background incorporation. **(B)** Ether phospholipids regulate choline incorporation strongly dependent on their ether-linked alkyl chain at sn-1 position and ester-linked acyl chain at sn-2 position. Either diacyl phospholipid (e.g. 16:0/18:0 PC) or dialkyl phospholipid (e.g. 16:0o/16:0o PC) prevents choline incorporation. Note that the polyunsaturated fatty acids (PUFAs) at sn-2 position may enhance choline incorporation (16:0/20:4 PC). **(B1)** Representative images of incorporated choline-O-Cy2 in WT and T2KO bEnd.3 cells with or without treatment with different phospholipids (PC-O, PC, or LPC). **(B2)** Statistical graphs of CTCF of the images. n = 12, ***p ≤ 0.001. Scale bar, 5 µm. **(C)** Dynamic analysis of choline uptake using live cell station revealed that TPP2 depletion drastically impairs this process (**bottom left**), which can be almost completely rescued by PC-O (**top right**) or E2 (**bottom left**) treatment. **(D)**TPP2 depletion significantly increases plasma membrane fluidity and metabolic dynamics and that PC-O (C18:0p/18:1 PC) treatment can strongly restore plasma membrane rigidity without altering incorporated ester-linked PC in plasma membrane. **(D1)** Transmission electron microscopic imaging revealed that the trilaminar or railroad track structure of the plasma membrane is interrupted in the absence of TPP2 and that PC-O treatment can significantly restore the trilaminar structure. The images below are enlarged versions of those above, presenting the regions of interest (ROI) in greater detail. Scale bar, 30 nm (upper), 10 nm (lower)**. (D2)** Steady-state fluorescence anisotropy using the gold-standard hydrophobic probe TMA-DPH (1-(4-trimethylammoniumphenyl)-6-phenyl-1,3,5-hexatriene) revealed that the surface fluidity of plasma membrane in the absence of TPP2 significantly increases and that PC-O treatment can almost completely restore the surface rigidity (upper). In addition, steady-state fluorescence anisotropy using the fluorescent PC analog BODIPY FL C5-HPC (1-Hexadecanoyl-2-(4,4-Difluoro-5,7-Dimethyl-4-Bora-3a,4a-Diaza-s-Indacene-3-Pentanoyl)-*sn*-Glycero-3-Phosphocholine) revealed that the plasma membrane metabolic dynamics is more active in the absence of TPP2 compared to WT control and that PC-O treatment doesn’t significantly change incorporated PC analog in plasma membrane (lower). **(E)**Choline uptake is affected in cerebrovascular ECs specifically upon FLVCR2 depletion. **(E1)** RT-PCR analysis of choline transporter expression in primary ECs and bEnd.3 cells demonstrated that FLVCR2, CTL1, and CTL2 are all expressed. **(E2)** Choline uptake assay revealed that depletion of FLVCR2 specifically impairs choline uptake in bEnd.3 cells, whereas depletion of CTL1 had no effect. Choline uptake was not assessed for CTL2 transporter, as it is normally localized to the mitochondrial membrane rather than the plasma membrane. Representative results from three independent experiments are shown. Scale bar: 5 µm. **(F)** Molecular docking simulation revealed that the choline transporter FLVCR2 and PC-O (C18:0p/18:1 PC) exhibit high binding affinity through hydrogen-bond and Pi-alkyl interaction. The docking score reaches -11.125525 kcal/mol. **(F1**) The three-dimensional schematic depicts the deep view of the interaction between FLVCR2 and PC-O. Key residues are shown in a ball-and-stick model, the protein FLVCR2 in a colored cartoon representation, and the ligand PC-O in gray. Nitrogen and oxygen atoms are colored blue and red, respectively. **(F2**) The molecular interaction is shown in a two-dimensional diagram. In this conformation, the PC-O ligand mainly contacts FLVCR2 through hydrogen bonds and Pi-alkyl interaction. Specifically, FLVCR2 ASP147 forms two hydrogen bonds with the glycerol backbone of PC-O, and FLVCR2 PHE347 forms Pi-alkyl bond with the ether linked alkyl chain of PC-O. **(G)** Both FLVCR2 ASP147ALA (D147A) and FLVCR2 PHE347ALA (F347A) mutations resulted in complete loss of choline transport function. **(G1)** DNA sequencing results of FLVCR2 D147A and FLVCR2 F347A mutations**. (G2)** Choline uptake **i**n FLVCR2-depleted bEnd.3 cells was rescued by ectopic expression of FLVCR2 but not by ectopic expression of its D147A and F347A mutants. Representive images from more than 100 images taken are shown. Scale bar, 10 µm.

With the fluorescent choline uptake assay established, we next performed a screening of various ether phospholipid species to assess their impact on choline uptake efficiency. Our criteria for lipid selection were based on a reference indicating that Far1 prefers saturated and unsaturated fatty acyl-CoAs of 16–18 carbon atoms as substrates^68^. Our results revealed that for ether phospholipids to effectively rescue the choline uptake deficiency caused by TPP2 depletion, an sn-1 ether linkage is essential, and any sn-2 substituent must be ester-linked (Fig. 6B). Both diacyl phospholipid (e.g. 16:0/18:0 PC) and dialkyl phospholipid (e.g. 16:0o/16:0o PC) prevent choline incorporation (Fig. 6B). Note that an ether linkage at the sn-1 position of glycerol backbone (C18:0p LPC) is sufficient to trigger high choline uptake, while the ester-linked PUFA at sn-2 (e.g., 16:0/20:4 PC) may further enhance choline incorporation (Fig. 6B). To corroborate the rescue effect of PC-O on choline uptake, we employed live-cell imaging to perform a dynamic analysis. Our results revealed that TPP2 depletion drastically impairs this process (Fig. 6C, bottom left), which can be almost completely rescued by either PC-O (Fig. 6C, top right) or E2 (Fig. 6C, bottom left) treatment.

To elucidate why the sn-1 ether linkage is essential for choline uptake, we investigated its impact on plasma membrane properties. First, TEM revealed that TPP2 depletion disrupts the characteristic trilaminar (“railroad track”) structure of the plasma membrane, and this structural integrity was significantly restored by PC-O (C18:0p/18:1 PC) treatment (Fig. 6D1). We then quantified membrane dynamics using steady-state fluorescence anisotropy. Using the hydrophobic probe TMA-DPH, which reports on surface fluidity, we found that TPP2 depletion significantly increases membrane fluidity, an effect that is nearly completely reversed by PC-O (Fig. 6D2, upper). In contrast, anisotropy measurements using the fluorescent PC analog BODIPY FL C5-HPC, which is commonly used in membrane dynamics studies, revealed more active membrane lipid metabolic dynamics in TPP2-depleted cells than in WT controls. However, PC-O treatment only produced a non-significant trend toward reducing the abundance of this PC analog in the membrane (Fig. 6D2, lower), suggesting that PC-O may actually increase membrane metabolic dynamics even while it significantly increases membrane rigidity.

We assumed that membrane properties may influence the stability of choline transporters. First, we examined the expression profiles of all three previously reported choline transporters (FLVCR2, CTL1, and CTL2) in primary brain vascular cells and bEnd.3 cells using RT-PCR and found that their expression levels are comparable between T2KO cells and WT controls (Fig. 6E1). Subsequently, immunoblot analysis of these transporters in plasma membrane or in mitochondria also revealed no significant differences between the T2KO cells and WT controls (data not shown), indicating that TPP2 depletion does not affect the expression levels of any of the three choline transporters, and that PC-O treatment does not rescue their choline uptake function in TPP2-depleted cells by protecting their stability. To further investigate the underlying mechanism, we generated FLVCR2- and CTL1-knockout bEnd.3 cell lines to assess their respective roles in choline uptake by cerebrovascular ECs. Since CTL2 is predominantly localized to the mitochondrial membrane rather than the plasma membrane^69^, corresponding knockout lines were not established for this protein. Our choline uptake assays conducted with the FLVCR2- and CTL1-knockout bEnd.3 and 293T cells demonstrated that choline uptake in either vascular ECs or 293T cells is specifically impaired upon FLVCR2 depletion (Fig. 6E2, 293T data not shown).

It was reported that FLVCR2 is a BBB (Blood Brain Barrier) choline transporter and is responsible for the majority of choline uptake into the brain^70^. To investigate how PC-O regulates the choline uptake function of FLVCR2, we further examined whether there is a physical interaction between FLVCR2 and PC-O. We first performed molecular docking simulation and found that the choline transporter FLVCR2 and PC-O (C18:0p/18:1 PC) exhibit high binding affinity through hydrogen-bond and Pi-alkyl interaction. Specifically, FLVCR2 ASP147 forms two hydrogen bonds with the glycerol backbone of PC-O, and FLVCR2 PHE347 forms Pi-alkyl bond with the ether-linked alkyl chain of PC-O. The docking score reaches - 11.125525 kcal/mol (Fig. 6F). To determine whether these interactions in situ are directly responsible for choline uptake, we generated two site-directed mutants. Crucially, both FLVCR2 ASP147ALA (D147A) and FLVCR2 PHE347ALA (F347A) mutations resulted in complete loss of choline transport function of FLVCR2 (Fig. 6G).

### AAV-mediated expression of Far1 in cerebrovascular ECs not only rescues memory impairment in female mice caused by TPP2 depletion, but also significantly ameliorates learning and memory deficits resulting from natural aging

In light of the critical role of PC-O in choline uptake—a process closely linked to cerebrovascular degeneration—we have generated an adeno-associated virus (AAV) vector for endothelial cell-specific expression of Far1 or CYP19A1. This approach enabled us to assess whether Far1-associated PC-O biogenesis and CYP19A1-associated estrogen production can ameliorate the cerebrovascular degeneration caused by TPP2 depletion. We performed the whole brain ex-vivo imaging and immunoblot and demonstrated that both Far1 and CYP19A1 fusion proteins are successfully expressed following intravenous delivery of HBAAV2/Br1-Tie1-Far1-Flag-ZsGreen and HBAAV2/Br1-Tie1-CYP19A1-Flag-T2A-ZsGreen viral particles. Interestingly, ectopic expression of Far1 fusion protein also moderately increases endogenous Far1 level (Fig. 7A). We then performed FDISCO imaging of the mouse hippocampal vascular system and demonstrated that either Far1 or CYP19A1 fusion protein expression significantly increases vascular volume and vascular diameter in both the CA1 and DG regions of female T2KO mouse brains (Fig. 7B). However, total vascular length of these mice showed a significant increase only in the CA1 region, while the increase observed in the DG region was not statistically significant (Fig. 7B).

**Fig. 7.**
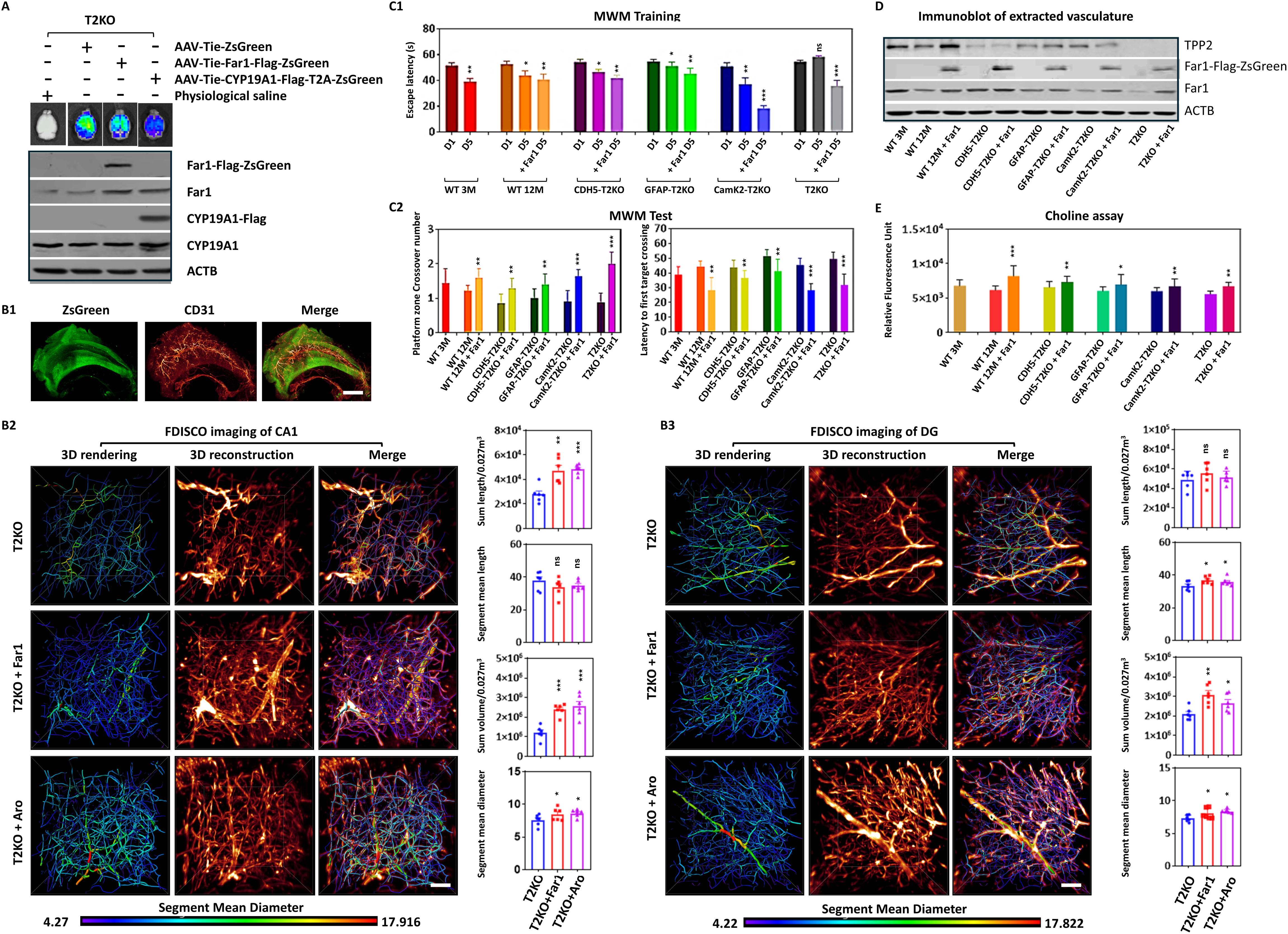
AAV-mediated expression of Far1 in cerebrovascular ECs not only rescues learning and memory impairment in female mice caused by TPP2 depletion, but also significantly ameliorates learning and memory deficits resulting from natural aging. **(A)** Whole brain ex-vivo imaging and immunoblot showed that Far1-Flag-ZsGreen and CYP19A1-Flag-T2A-ZsGreen are expressed after intravenous delivery of HBAAV2/Br1-Tie1-Far1-Flag-ZsGreen and HBAAV2/Br1-Tie1-CYP19A1-Flag-T2A-ZsGreen virus particles. The endogenous Far1 and CYP19A1 served as an internal control. **(B)** FDISCO imaging of the mouse hippocampal vascular system revealed that either Far1-Flag-ZsGreen or CYP19A1-Flag expression significantly increases vascular volume and vascular diameter in both the CA1 and DG regions. However, total vascular length showed a significant increase only in the CA1 region, while the increase observed in the DG region was not statistically significant. **(B1)** Representative FDISCO image of the mouse hippocampus. ZsGreen fusion protein (**green**) and blood vessels stained for CD31 (**red**) are shown. Morphometric analysis of the FDISCO images of CA1 regions **(B2)** and DG regions **(B3)** by 3D reconstruction and rendering. Representative 3D reconstructions are shown on the left; corresponding quantitative graphs for vascular total length, mean segment length, total volume, and mean diameter are on the right. The segment mean diameter is represented by a color gradient, increasing from blue to red. Scale bar, 50 µm. **(C)** Morris water maze test showed that intravenous delivery of HBAAV2/Br1-Tie1-Far1-Flag-ZsGreen virus significantly rescues the learning and memory impairment of female global and conditional T2KO mice. **(C1)** Analysis of escape latency (Day 1 vs. Day 5 of training) showed that global T2KO mice failed to learn, whereas conditional T2KO mice showed significantly impaired learning and memory. However, injection of HBAAV2/Br1-Tie1-Far1-Flag-ZsGreen virus particles significantly rescued learning and memory deficits in all knockout mice. **(C2)** A comparison of the number of platform crossings and escape latency during the test phase demonstrated that above-mentioned viral injection significantly improves learning and memory ability in all groups of knockout mice. ***p ≤ 0.001, **p ≤ 0.01, *p ≤ 0.05; WT 3M (3 month-old WT), n = 11; WT 12M (12 month-old WT), n = 18; WT 12M + Far1, n = 10; CDH5-T2KO, n = 14; CDH5-T2KO + Far1, n = 10; GFAP-T2KO, n = 14; GFAP-T2KO + Far1, n = 10; CamK2-T2KO, n = 8; CamK2-T2KO + Far1, n = 6; T2KO, n = 21; T2KO + Far1, n = 6. **(D)** Immunoblot analysis of isolated vasculature revealed that viral delivery not only induces successful ectopic Far1 expression but also further elevate the levels of endogenous Far1 (Fig. 7D). It is noteworthy that both endogenous TPP2 and Far1 abundance are significantly reduced in isolated vasculatures of 12-month-old female mice compared to that of 3-month-old female mice. Of particular importance, ectopic expression of Far1 markedly increases TPP2 level in vasculature of 12-month-old naturally aged mice. **(E)** Choline measurements indicated that ectopic Far1 expression following viral delivery significantly elevates cerebral choline content in the mouse brains. ***p ≤ 0.001, **p ≤ 0.01, *p ≤ 0.05, n = 12.

**Fig. 8.**
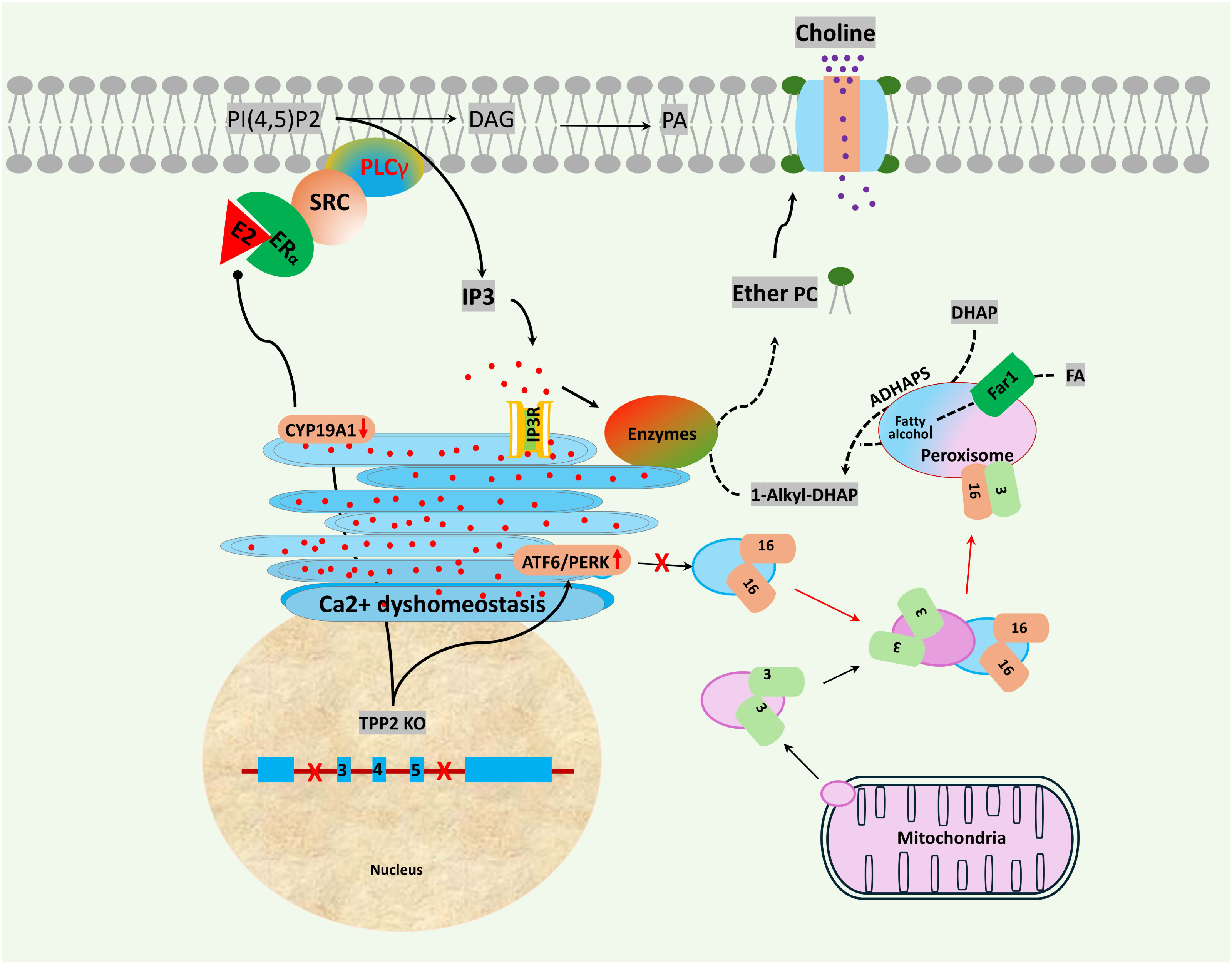

Encouraged by the FDISCO imaging results, we conducted the Morris water maze (MWM) test to assess whether ectopic expression of Far1 could enhance the learning and memory abilities of global female T2KO mice, concomitant with improvements in vascular density and overall vascular health. Our findings demonstrated that ectopic expression of Far1 not only restores learning and memory abilities in global female T2KO mice, but also significantly alleviates learning and memory impairments in conditional T2KO mice (Fig. 7C). In more details, during the training phase, global T2KO mice exhibited a complete failure to learn, while conditional T2KO mice displayed significantly impaired learning and memory. However, ectopic expression of Far1 substantially restored the learning and memory deficit in all T2KO mice (Fig. 7C1). In the test phase, comparisons of platform crossings and escape latency further confirmed that ectopic expression of Far1 significantly improved learning and memory performance in both global and conditional T2KO mice (Fig. 7C2). Of particular importance, administration of Far1 expressing virus also significantly enhanced learning and memory performance in naturally aged mice (Fig. 7C).

To confirm the ectopic expression of Far1 after administering the Far1 expressing virus particles, as well as its effect on elevating brain choline levels, we conducted immunoblot analysis and choline assays. Our immunoblot analysis of isolated vasculature revealed that viral delivery not only induced successful ectopic Far1 expression but also further elevated the levels of endogenous Far1 (Fig. 7D). It is noteworthy that both endogenous TPP2 and Far1 abundance are significantly reduced in isolated vasculatures of 12-month-old female mice compared to that of 3-month-old female mice (Fig. 7D). Consistent with this finding, choline measurements indicated that ectopic Far1 expression following viral delivery significantly elevates cerebral choline content in the mouse brains of all T2KO mice and naturally aged mice (Fig 7E).

## DISCUSSION

This study provides the first evidence that TPP2 is critical for maintaining cerebrovascular homeostasis in adult female mice (Fig. 1A-G). Mechanistically, TPP2 orchestrates proper intracellular Ca²⁺ distribution (Fig. 4A-C) and local CYP19A1/estrogen homeostasis in cerebrovascular ECs and perivascular astrocytes (Fig. 4D2, E, F, I, and Suppl. Fig. 3A-B). Specifically, TPP2 deficiency disrupts the anticipatory UPR signaling (Fig. 4G-H) and consequently drives aberrant pexophagic flux (Fig. 5A-F), which depletes PC-O (Fig. 3A-B) and further causes subsequent choline depletion (Fig. 2A, C-E) as well as significantly decreased ACh (Fig. 2B and F). It is noteworthy that we employed the bEnd.3 cell line, which is derived from mouse brain endothelium, for some of experiments to complement our *in vivo* T2KO models and primary cerebrovascular ECs. Although the sex of the bEnd.3 cell line is not documented in available sources, we confirmed its robust CYP19A1 expression, validating its utility for our experiments (Fig. 4D2).

Our work in neurons revealed that TPP2 sustains CYP19A1 levels by maintaining Ca²⁺ homeostasis and PC biosynthesis. Without this regulation, disrupted Ca ²⁺ homeostasis and PC biosynthesis could otherwise induce ER stress and subsequently trigger CYP19A1 degradation via ER stress-associated autophagy^71^. We demonstrate in this study that the anticipatory UPR signaling pathway, which constitutes a critical nexus between cytosolic estrogen levels and homeostatic Ca²⁺ distribution, is nearly completely disrupted in vascular ECs in the absence of TPP2 (Fig. 4G). Furthermore, consistent with findings in neurons, TPP2 physically interacts with IP3R1, SERCA1, and VDAC2 in vascular ECs (Fig. 4D1). However, unlike in neurons where IP3R1 levels remain stable, vascular ECs exhibit a significant decrease in IP3R1 (Fig. 4D2). These findings indicate that physical interaction with TPP2 is crucial for maintaining IP3R1 stability in cerebrovascular ECs. This notion was corroborated by the finding that ectopic expression of catalytically inactive TPP2 mutants (TPP2 S449A and S449T) specifically rescued the IP3R1 deficiency in T2KO cells (Supplementary Fig. 3C), highlighting a potential structural or scaffolding role for TPP2 in this context. This finding suggests that the function of a given molecule such as IP3R1 can be modulated in a cell-type-specific manner. Collectively, we inferred that in the absence of TPP2, the combined deficiency in anticipatory UPR signaling and IP3R1 could lead to severe Ca²⁺ dyshomeostasis and a nearly complete loss of CYP19A1/estrogen in vascular ECs. This was indeed the case. Our two-photon imaging clearly showed that almost no Ca²⁺ signal was detected by Fluo-4 staining in T2KO hippocampal vascular cells (Fig. 4A), while immunofluorescence staining revealed significantly reduced levels of CYP19A1 in hippocampal vascular ECs of T2KO 12-month-old mice compared to WT controls (Fig. 4E). Consistently, targeted metabolomic results fully confirmed that estrogen is nearly undetectable within these cells of T2KO 12-month-old mice compared to WT controls (Fig. 4F). Moreover, there is a precise synchrony between Ca²⁺ imbalance and CYP19A1 depletion, and restoring Ca²⁺ homeostasis can concomitantly ameliorate CYP19A1 deficiency (Fig. 4I). Additionally, inhibition of autophagic flux with 3-MA rescues the CYP19A1 depletion caused by TPP2 loss in vascular ECs (Fig. 4I), indicating that TPP2 maintains CYP19A1 abundance by suppressing an autophagic pathway triggered by Ca²⁺ imbalance. In summary, TPP2 plays a critical role in maintaining intracellular Ca²⁺ distribution and local estrogen levels in the adult female vasculature, while acknowledging that the underlying mechanism differs a little bit from that observed in neurons.

Ether lipids have been shown to be involved in a variety of biological processes. In particular, the influence of ether phospholipids on cell membrane structure and fluidity is profound and multifaceted, primarily due to their unique chemical architecture^72^. This study provides the first evidence that TPP2 is essential for maintaining ether-PC levels by sustaining peroxisomal homeostasis (Fig. 5A-F). Ether phospholipids (especially plasmalogen of 16–18 carbon atoms) are essential for choline uptake across brain vascular ECs (Fig. 6B and 6C). Therefore, we examined whether ether phospholipids affect choline uptake by altering plasma membrane fluidity and dynamics. Our results demonstrate that while ether phospholipids significantly increase membrane rigidity in T2KO cells, they do not markedly decrease plasma membrane dynamics (Fig. 6D1 and 6D2). In fact, membrane dynamics even appear to be slightly enhanced after treatment with ether phospholipid (Fig. 6D2 lower). To further explore how ether phospholipids affect choline uptake independent of their influence on cell membrane dynamics, we performed molecular docking simulations. We postulated that ether phospholipids may regulate choline transporter function through direct or indirect physical regulation. Our in silico research results revealed that a typical species of ether phospholipids interacts with FLVCR2 with very high affinity (docking score: -11.125525 kcal/mol) through forming hydrogen bonds with FLVCR2 D147 and Pi-alkyl bond with FLVCR2 F347. To validate the accuracy of this prediction, we generated the site-directed mutants of these two amino acid residues to determine whether these two ether phospholipids interacting sites are indeed critical for choline uptake through FLVCR2. Astonishingly, any one of the two site mutations completely abolishes choline uptake function of FLVCR2 (Fig. 6G), implicating that ether phospholipids most likely regulate FLVCR2 function allosterically by physically interacting with these two residues. Our findings open a new avenue for the artificial regulation of choline intake, offering boundless possibilities for the treatment of diseases related to choline deficiency.

Choline is an essential nutrient that serves as a precursor for several critical molecules in the nervous system, most notably the neurotransmitter ACh and the membrane PC. Adequate choline intake is therefore crucial for maintaining normal brain function. Dysfunctions in choline intake and choline metabolism may drive the development or advancement of several neurological disorders, such as AD, mild cognitive impairment (MCI), and Mood disorders^73–76^. In this study, we demonstrate that choline starvation causes severe cerebrovascular degeneration as observed in the hippocampal region of female T2KO mice (Fig. 2H). As abovementioned, TPP2 plays a crucial role in maintaining brain choline and ACh homeostasis by modulating choline uptake across brain vascular ECs (Fig. 2A-2E). Mechanistically, TPP2 regulates choline uptake by modulating ether-PC levels, which may allosterically regulate the choline uptake function of FLVCR2 (Fig. 3B-3E and Fig. 6). Based on these findings, we infer that TPP2 deficiency initiates a vicious cycle in which impaired ether phospholipid synthesis and compromised choline uptake exacerbate each other, thereby leading to a severe depletion of PC-O and ACh and culminating in vascular degeneration. Indeed, this study showed that TPP2 depletion leads to cerebrovascular degeneration characterized by decreased EC number and their mislocalization within narrowed vascular lumens (Fig. 1A, C-E). More strikingly, our scRNA-seq data revealed an approximately 50% reduction in the EC population, as defined by a five-gene signature (Fig. 1F-G). However, dynamic contrast-enhanced MRI revealed no evidence of increased blood–brain barrier permeability or vasogenic edema in T2KO mouse brains, despite the observed vascular degeneration (Fig. 2G). These seemingly contradictory observations can be attributed to two possible explanations. One is that TPP2 depletion does not substantially disrupt the overall connectivity of ECs, which form tight junctions with one another, despite the altered expression of their signature genes resulting from estrogen deficiency. Consistent with this notion, three of the five signature genes (CD34, Kdr, and Cldn5) are known to be estrogen-regulated. The other one is that TPP2 depletion leads to deficiency of PC and ACh (a master regulator of vascular dilation) as abovementioned, thereby narrowing the blood vessel lumen and compressing ECs together.

Cebrovascular degeneration plays a significant role in the pathogenesis of both VaD and AD^3, 13^. Although estrogen serves as a potent endogenous protector of the vascular system in females throughout their lifespan^77^, ERT/MHT is currently not approved for use in preventing or treating VaD and AD due to its adverse effects on other organs and the stringent requirement for precise timing of treatment^78^. It is now solely confined to the management of menopausal symptoms^54^. Thus, new therapeutic strategies are required to maximize protective effects of estrogen within the brain while minimizing peripheral risks and overcoming the limitations of the treatment window. In fact, new therapeutic strategies, particularly those involving SERMs^55–59^ and approaches aimed at boosting or restoring local neurosteroidogenesis within the brain^60–62^ are being explored. However, to our knowledge, although several candidate drugs related to this aim have been developed, they are currently still in the stage of clinical trials or transitioning from preclinical research to clinical application^56, 57, 61, 62^. While tibolone is employed clinically, concerns remain due to reported increases in the relative risk of hormone-sensitive cancers^55^. Our findings reveal that TPP2 sustains cerebrovascular integrity by orchestrating a series of interconnected events: proper intracellular Ca²⁺ distribution, local estrogen biosynthesis, ethe-PC synthesis, and ACh production. Building on this mechanism, a novel strategy targeting TPP2 and its interconnected effectors to prevent cerebrovascular degeneration may precisely address this therapeutic need in both VaD and AD.

Consistent with its potential as a therapeutic target for VaD and AD, we observed that endogenous TPP2 levels in isolated brain vasculature were markedly decreased in 12-month-old female mice compared to 3-month-old controls (Fig. 7D). Although 12-month-old mice do not yet exhibit overt clinical phenotypes of AD or VaD, they do show an obvious decline in learning and memory capabilities compared to 3-month-old mice (Fig. 7C). Together with our findings in T2KO mice, these observations suggest that the age-dependent decline in TPP2 vascular expression may play a key role in age-related memory impairment. Therefore, a key therapeutic objective needs to identify the mechanism that sustains vascular TPP2 abundance and function into later life. Apropos to this requirement, we found that TPP2 deficiency in naturally aged mice can be restored by AAV-mediated ectopic expression of Far1 in vascular ECs (Fig. 7D). Regarding the underlying mechanism, which remains a subject for future investigation, we hypothesize that Far1 regulates TPP2 levels via its established function in ether phospholipid biosynthesis. It is noteworthy that TPP2 depletion, in turn, markedly downregulates Far1 levels in peroxisomes (Fig. 5B and 5C). Considering that TPP2 depletion induces estrogen deficit (Fig. 4F), which further leads to severe peroxisome deficit (Fig. 5A) and results in inadequate ether phospholipid production (Fig. 3B), we assume that Far1 and TPP2 may form a mutually regulatory feedback loop, potentially via estrogen and ether phospholipid, both of which have been shown to be closely related (Fig. 3B2 and B5) and play crucial roles in preserving vascular health. Indeed, AAV-mediated ectopic expression of either Far1 (for PC-O biosynthesis) or CYP19A1 (for estrogen biosynthesis) significantly increases vascular volume and vascular diameter in both the CA1 and DG regions (Fig. 7B). Of note, total vascular length showed a significant increase only in the CA1 region, while the increase observed in the DG region was not statistically significant following injection of Far1 expressing virus particles (Fig. 7B). Regarding the regional differences in effects of estrogen and ether phospholipids on blood vessels, the underlying mechanism is unclear. Although the hippocampal CA1 and DG regions are adjacent, they exhibit significant differences in cellular composition, vascular networks, and function^59, 79^. The effects of estrogen and ether phospholipid are likely to vary due to the distinct local microenvironments of these different regions. Nonetheless, AAV-mediated ectopic expression of Far1 rescues memory impairment caused by both global and conditional TPP2 depletion (Fig. 7C). Of particular importance, it also significantly ameliorates learning and memory decline resulting from natural aging (Fig. 7C).

In summary, this study has revealed a novel mechanism underlying brain vascular degeneration in adult female mice. Specifically, over time, TPP2 levels in brain vascular cells decrease significantly, triggering a cascade of pathological events, namely Ca²⁺ imbalance, estrogen depletion, aberrant pexophagy, impaired ether phospholipid biosynthesis, and dysregulated choline uptake and ACh production. Within this signaling pathway, two vicious cycles are established: one between Ca²⁺ imbalance and estrogen deficiency, and the other between ether phospholipid deficit and choline uptake dysregulation. Together, these processes ultimately drive brain vascular degeneration. To counteract the vascular degenerative progression, we developed an AAV-mediated therapeutic strategy designed to either prevent natural aging-associated memory decline or treat VaD and AD disorders.

## MATERIALS AND METHODS

### Animals and antibodies

Mice with conditional knockout of the TPP2 gene in ECs, NSCs, and astrocytes were prepared by hybridizing floxed TPP2 mice with CDH5-Cre (Shanghai model organisms, Shanghai, China), Nestin-Cre, and GFAP-Cre mouse lines (Cyagen, Suzhou, China), respectively. Mice with conditional knockout of TPP2 in microglia were obtained by crossing floxed TPP2 mice with TREM119-2A-CreERT2 mouse line (Cyagen, Suzhou, China) combined with injection of Tamoxifen (MCE, Shanghai, China). Global T2KO mice (Suppl. Fig. 1) and floxed mouse line (Suppl. Fig. 2) used in this project are as described previously^71^. In addition, Detailed information regarding the antibodies used in this study, including the manufacturers, working concentrations, and experimental applications, is provided in the antibody list (Suppl. table 1).

### Cell lines and plasmids

The bEnd.3 and 293T cell lines were obtained from ATCC (Gaithersburg, USA). For the generation of gene knockout bEnd.3 and 293T cell lines, all-in-one CRISPR/Cas9 plasmids were constructed by inserting sgRNAs downstream of the U6 promoter in either the YKO-LV010 (Ubigene bioscience, Guangzhou, China) or pPB[CRISPR]-Hygro-hCas9-U6 vector (Hysigen Bioscience, Suzhou, China). The murine sgRNA sequences are as follows: Tpp2 (sgRNA: 5’-CGTATTGAGAAACTACAAAG-3’), Atf6 (sgRNA: 5’-TCGACGTTGTTTGCTGAACT-3’), Ern2 (sgRNA: 5’-TGTCCCGAGCCAGTGTGAGA-3’), Eif2ak3 (5’-CACTTCTCACTGCCGCTTCG-3’), Marchf5 (sgRNA:5’-AGATGATAGAACAGCTGAGT-3’), Chip (sgRNA: 5’-TGCTGACTGCCGGCGAGCCC-3’), Flvcr2 (5’-TCGGACAATGGAACTTAAGT-3’), Slc44a1 (5’-CTGGAAGCAATACCGAACAG-3’). The human sgRNA sequences are as follows: Flvcr2 (5’-TGATCTGGCACTACCCGGTA-3’), and Slc44a1 (5’-GCAAGTCTACTCACTTCCGC -3’). Gene knockout and double knockout cell lines were established according to standard protocol. In brief, cells were transfected with the genome engineering plasmid and were selected for about 2 weeks in complete medium (DMEM medium supplemented with 10% fetal calf serum and antibiotics) containing 1.2 µg/ml puromycin (Solarbio, Beijing, China) or 200 µg/ml hygromycin (Roche, Basel, Switzerland). Then gene knockout individual clones were isolated, expanded and identified by western blot and antibody staining. Plasmids for transient expression of Flvcr2 and Slc44a1 and their mutants were constructed by help of Hysigen Bioscience (Suzhou, China). Plasmids for transient expression of Tpp2 and its S449A mutant were constructed by help of SinoBiological (Beijing, China). Other plasmids for transient expression including pCAGGS-GCaMP2, pCMV G-CEPIA1er, pmScarlet-I_peroxisome_C1, pRRLSIN.cPPT.PGK-GFP-PTS1.WPRE were from Addgene (Watertown, USA).

### Chemical reagents and lipids

Chemical reagents including E2, PP2, CCT129957, 3-MA, CND1163, and D-myo-inositol-1,2,4,5-tetrakisphosphate tetrasodium salt (IP4 tetrasodium), C18:0p LPC, C18:0p/18:1 PC, 18:0p/20:4 PC, 18:0p/20:4 PE, 16:0/18:0 PC, 18:0/20:4 PC, 16:0p/16:0 PC, 16:0o/16:0o PC were obtained from MCE (Shanghai, China). Thapsigargin (TG) was obtained from GLPBIO (Montclair, USA). 1-(1Z-octadecenyl)-2-oleoyl-sn-glycero-3-phosphocholine (C18:0p/18:1 PC) was obtained from ZZBio (Shanghai, China). All reagents were used in accordance with the manufacturers’ instructions. In brief, the treating conditions were at 37°C combined with following concentrations and durations, namely, 50 nM E2 for 12h, 5 µM TG for 5 h, 5 µM C18:0p/18:1 PC, C18:0p LPC, 18:0p/20:4 PC, 18:0p/20:4 PE, 16:0/18:0 PC, 18:0/20:4 PC, 16:0p/16:0 PC, 16:0o/16:0o PC for 24 h, 10 µM PP2 for 24 h, 15 µM CCT129957 for 24 h, 2 mM 3-MA for 5 h, 10 µM CDN1163 for 24 h, 200 nM IP4 for 24 h.

### ACh and choline assay

Total ACh was quantified using Ach Elisa kit (Sangon Biotech, Shanghai, China) according to the manufacturer’s instruction. In brief, a total of 20 mg of brain tissue per sample was collected and rinsed once with cold 1 × PBS, and was homogenized in 200 µL 1 × PBS containing protease inhibitors. After centrifugation, 50 µL of each sample was used for ACh quantification. The subsequent protocol involving incubation with biotin-conjugated ACh antibody followed by HRP-conjugated streptavidin incubation, and substrate reaction was performed according to the manufacturer’s instructions. Absorbance was then measured at 450 nm using a microplate reader for quantification. Total choline levels were quantified using a commercial detection kit (abcam, Cambridge, USA) according to the manufacturer’s protocol. Briefly, 5 × 10^6^ cells or 20 mg of tissue were harvested, washed once with cold 1× PBS, and homogenized in 400 µL assay buffer. Following centrifugation at 4°C, the supernatant was transferred to a new tube and kept on ice. The choline reaction mixture was prepared immediately before use by adding 20 µL of 250× Red Dye stock solution (dissolved in 40 µL DMSO) to the choline probe (dissolved in 5 ml assay buffer). Then, 50 µL of the reaction mixture and 50 µL sample were added to each well (total volume 100 µL/well). The plate was protected from light and incubated at room temperature for 30 minutes. Fluorescence was monitored using a microplate reader at Ex/Em = 540/590 nm (with a cutoff at 570 nm).

### MOST imaging of mouse cerebrovascular atlas

Mice to be tested were fasted overnight and then anesthetized with an intraperitoneal injection of Zoletil® 50 (Virbac, Carros, France). The mice were subsequently perfused with 0.01M PBS followed by 4% paraformaldehyde fix solution. After perfusion, the intact brains were carefully extracted from the skulls and postfixed in 4% paraformaldehyde for 24 h at room temperature. The whole-mount Nissl staining method was as previously described^80^. In brief, brain samples were rinsed in 0.01 M PBS for 24 h, and then were stained for at least 12 days in a 2.5% thionine (MERCK, Darmstadt, Germany) solution. The stained brains were dehydrated firstly in an increasing gradient of ethanol (50%-70%-85%-95%-100%, every change 2 h), and followed in a 1:1 ratio of 100% ethanol and 100% acetone for 2 h, 100% acetone for 2 h, 100% acetone overnight, and another 100% acetone for an extra 24 h. Subsequently, samples were infiltrated with a graded series of Spurr resin (SPI supplies, West Chester, USA) in acetone (50%-75%-100% Spurr resin, every change 12 h), then 100% Spurr solution for 2 X 1.5 days. All the steps were performed at room temperature under rocking conditions. At last, the brains were placed in a silicone mold filled with 100% Spurr resin and then polymerized for 36 h at 60°C. The intact resin-embedded mouse brains were sectioned and imaged continuously for about 190 hours with a sectioning thickness of 1.5 μm. The image tile was obtained with a voxel size of 0.35 μm × 0.35 μm × 1.5 μm, using a 40× objective and a time-delay integration (TDI) line-scan charge-coupled device^81^. Each brain included more than 8000 coronal sections. The raw images were preprocessed for seamless image stitching and luminance nonuniformity correction^82^. After that, an image optimization technique was used to further improve background uniformity, periodic noise, and to enhance the image contrast of the preprocessed coronal images. Several steps including background correction, noise reduction, and contrast enhancement were implemented to perform image optimization^83^. The enhanced contrast of the optimized coronal images significantly improved the convenience of information extraction and grayscale rendering. The vascular vessels of the whole brain were extracted based on region growth algorithm using Amira 3D software (Thermo Fisher Scientific, Waltham, USA). The coronal and horizontal view of the whole brain vascular distribution was reconstructed based on the 300 μm-thick coronal sections from a representative brain of both the WT and T2KO mice. The regions of interest including cortex and hippocampus were also extracted to conduct morphological and quantitative comparison between the WT and T2KO mice. The main morphological parameters in the vascular assessment include the segment length, length density, volume fraction and diameter. The segment is defined as compartment lying between two branching nodes. The segment length is the length of the segment. The length density was used for measuring the total vascular length per imaging volume (meters per cubic millimeters). The volume fraction was adopted to calculate the ratio of the total vascular volume to the volume of the corresponding brain tissue.

### T1-weighted contrast-enhanced MRI

Mice for test were injected with 0.1 mL gadoteric acid meglumine salt solution (0.377g/mL, Jiangsu Hengrui Pharmaceutical Co., Ltd, China) via tail vein, and after waiting for 15 minutes and subsequent anesthesia with isoflurane (1.5-2% in O2) in Summit Anesthesia support (Patterson Scientific, Saint Paul, U.S.) T1-weighted images were acquired (repetition time 225.891 ms, echo time 3 ms, slice thickness 0.5 mm, slices 20, averages 4, image size 384 x 384, field of view 25 x 25 mm²) using BioSpec 94/20 (Bruker, Billerica, USA), with the animals being placed in a dedicated cylindrical cradle for reproducible positioning of the mouse head and the respiration rate being continuously monitored through the PC-SAM software interface (SA Instrument, NY, USA).

### Dynamic ^11^C-choline PET/CT

Mice for test were injected with 18.7 MBq N-[^11^C]methyl choline chloride (Shanghai Institute of Materia Medica, Shanghai, China) via tail vein, and subsequently anesthesized with isoflurane (1.5-2% in O2) in Summit Anesthesia support (Patterson Scientific, Saint Paul, USA). PET/CT scans were performed on Inveon 3D PET/CT scanners (SIEMENS, Munich, Germany) continuously for 60 min and images were acquired at 15 minute intervals with final anatomical localization by CT combined with CT-based Attenuation Correction (CTAC). In more details, for acquisition of PET/CT images the following parameters were used. For PET: Axial scan length-127 mm, lower level discrimination-350 keV, upper level discrimination-650 keV, Timing window-3.488 ns; and for CTAC CT: Total rotation-half, Projections-120, Settle time-0 ms, Magnification-low, Binning-4x4, Transaxial CCD size-3072 px, Axial scanning length-133 mm, Voltage-80 kV, Current-500 µA, Exposure-150 ms.

### Two-photon imaging of Ca2+ in brain slices

Acute transverse hippocampal slices (200 μm thick) were prepared from mouse brains using a vibratome (Leica Biosystems, Wetzlar, Germany), starting approximately 10 mm from the rostral end. Slicing was performed in ice-cold, oxygenated, modified artificial cerebrospinal fluid (ACSF) containing (in mM): 95 NaCl, 1.8 KCl, 1.2 NaH₂PO₄, 7 MgSO₄, 26 NaHCO₃, 15 glucose, and 50.5 sucrose. The slices were then transferred to a chamber and incubated for 30 min at room temperature in continuously oxygenated standard ACSF supplemented with 2 μM Fluo-4 AM (Beyotime Biotechnology, Shanghai, China). Ca²⁺ imaging in hippocampal neurons was performed using a two-photon microscope (Femto3D Atlas, Femtonics, Budapest, Hungary) equipped with a 25× water-immersion objective (MRD77220, NA 1.10, WD 2 mm; Nikon, Tokyo, Japan). Fluorescence excitation was achieved using a tunable laser set to a wavelength of 920 nm, and images were acquired at a repetition rate of 1.7 Hz.

### Spatial metabolomics analysis of the hippocampus

Mice for test were anesthesized and transcardially perfused with saline. Subsequently, the hippocampi were dissected and embedded in Cryo-Gel (Leica Biosystems, Nussloch, Germany). The samples were frozen at -80 °C, and then 10 μm-thick sagittal slices were prepared using a Cryostat Microtome (Leica Microsystems, Wetzlar, Germany) and were thaw-mounted on SuperfrostTM PLUS slides (Thermo Fisher Scientific, Waltham, USA). At least 4 pieces of serial slices for each hippocampus were prepared, one for H&E staining, three for mass spectrometry imaging (MSI). For MSI, sections were firstly frozen at -80 °C, and then desiccated in succession at -20 °C for 1 h and at room temperature for 2 h. Subsequently, MSI of the prepared sections were performed on a Desorption Electrospray Ionization (DESI) platform (Waters, Milford, USA) in conjunction with SYNAPT XS Mass Spectrometer (MS) (Waters, Milford, USA) as previously reported^84^. More specifically, the surface of the sample sections was constantly scanned according to the following parameters: the scanning rate in x direction-0.2 mm/s, the resolution in y direction-50 μm, the flow rate of acetonitrile/H2O (8:2, 0.1% formic acid)-1.5 μL/min, the transporting gas flow rate-45 L/min, the spray voltage-7 kV, the distance between sample surface and sprayer-3 mm, the distance from sprayer to ion transporting tube-3 mm. The analyzer scanned over a mass range of 70-1000 m/z at a resolution-70,000, the automated gain control target level-2e6, the maximum injection time-200 ms, the S-lens voltage-55 V, and the capillary temperature-350 °C. For data processing the collected raw files were converted into imzML format using imzMLConverter and then imported into MSiReader for ion image reconstructions after background subtraction using the Cardinal software package. All MS images were normalized using total ion count normalization (TIC) in each pixel. Region-specific MS profiles were precisely extracted by matching high-spatial resolution H&E images. The discriminating endogenous molecules of different tissue microregions were screened by a supervised statistical analytical method: OPLS-DA. Variable Importance of Projection (VIP) values obtained from the OPLS-DA model were used to rank the overall contribution of each variable to group discrimination. The VIP value reflects the importance degree on the classification of sample categories with respect to the first two principal components of the OPLS-DA model, which indicates that this variable has a significant effect if the VIP is greater than 1. A two-tailed Student’s T-test was further used to verify whether the metabolites of difference between groups were significant. Differential metabolites were selected with VIP values greater than 1.0 and p-values less than 0.05. Additionally, for the special data structure obtained from the MSI analysis, we also performed T-distributed stochastic neighbor embedding (t-SNE) and uniform manifold approximation and projection for dimension reduction (UMAP) on the MS data in each pixels for dimensionality reduction, respectively. The Spatial shrunken centroids clustering (SSCC) was applied for MSI data clustering to separate the sample based on the differences abundance of ions in each pixels. For analyte identification the ions detected by AFADESI were annotated by the pySM pipeline and an in-house SmetDB database (Lumingbio, Shanghai, China).

### Mass spectrometry-based lipidomics

To identify the variation of lipid species and the related effect of E2, the lipidomics analyses of T2KO bEnd.3 cells and their control cells were performed as follows. In short, the sample lipids were prepared as reported^85, 86^. Chromatographic separations were then performed on an ExionLC^TM^ AD system (SCIEX, Framingham, USA) equipped with Accucore^TM^ C30 column (100 × 2.1 mm, 2.6 µm, Thermo Fisher Scientific, Waltham, USA), which was maintained at 45℃. After column equilibration, 2μL of each sample was injected and the analytes were eluted with a gradient of acetonitrile/isopropanol prepared in 0.1% formic acid and 10 mM ammonium formate at a flow rate of 0.35 mL/min over 20 min^87^. The QTRAP^TM^ 6500+ LC-MS/MS system equipped with Turbo Ion Source (SCIEX, Framingham, USA) was used for data acquisition with the following parameters: Temperature 500℃; Ion spray voltage 5.5 kV for positive mode and -4.5 kV for negative mode; Gas 1, gas 2, and curtain gas at 45, 55, and 35 psi, respectively. The Multiple Reaction Monitoring (MRM) approach was used for mass spectrometry quantification, while the Linear Ion Trap (LIT) analyzer in the system was used for qualitative scan. Details on the LC-MS/MS method using QTRAP system can be found on SCIEX community. MultiQuant software was used for processing of large batches of data and efficient quantification of detected lipid species. The MetaboAnalystR package was used for multivariate statistical analysis (PCA, PLS-DA, and OPLS-DA). To screen for differential lipids ANOVA was used with the conditions as follows: VIP≥1, P value ≤ 0.05, change ratio ≥ 2 or ratio ≤ 0.5.

### TLC

Commercially pre-coated silica gel plates (Gugent, Hongkong, China) with loading strips were first immersed in a 0.15 M (NH₄)₂SO₄ solution for 30 seconds and subsequently allowed to activate in a covered container for two days. On the day of analysis, the plates were dried by baking at 120°C for 2.5 hours. Lipid samples, extracted from 50 mg of WT or T2KO bEnd.3 cells using 600 µL of a methanol : chloroform : formic acid (20 : 10 : 1) solvent mixture plus 300 µL 0.2 M H_3_PO_4_ and 1M KCl, along with standard compounds, were spotted onto the baseline of the activated TLC plates (1.5 cm from the bottom edge). The spotted plates were then placed vertically in a sealed chromatography tank containing a shallow layer of the developing solvent (acetone : toluene : water, 90 : 30 : 7), ensuring the solvent level remained below the applied spots. The solvent was allowed to ascend until the front nearly reached the top of the plate. After development, the plates were removed from the tank and left in a fume hood for the solvent to evaporate completely. To visualize the separated lipid bands, the plates were evenly sprayed with a 50% (v/v) sulfuric acid solution in water and then carefully heated on a hot plate. Lipids appeared as charred brown or black spots. The retention factor (Rf value) for each spot was calculated as the distance traveled by the spot divided by the distance traveled by the solvent front. Preliminary identification of lipids was achieved by comparing both Rf values and spot patterns against those of known standards.

### TEM

For hippocampal vascular analysis, mice were humanely euthanized via cervical dislocation followed by immediate craniotomy. The tissue blocks (1 mm³) from CA1 hippocampal subregion was microdissected and rapidly immersed in primary fixative containing 2.5% glutaraldehyde/4% paraformaldehyde in 0.1 M phosphate buffer (PB, pH 7.4). Concurrently, bEnd.3 murine brain ECs (wild-type vs. T2KO) were cultured to 80% confluence in DMEM complete medium. Following trypsinization (0.25% EDTA-trypsin), cell pellets were fixed in identical glutaraldehyde-containing buffer for 12 h at 4 °C. Pre-fixed cellular aggregates were subsequently encapsulated in 1% low-melting agarose (Sigma-Aldrich, Saint Louis, USA) for structural stabilization. All specimens underwent secondary fixation with 1% osmium tetroxide (OsO₄, Ted Pella, Redding, USA) in PB for 2 h at 4 °C, followed by progressive ethanol dehydration series (30%, 50%, 70%, 80%, 95%, 100% X 2) with 20 min intervals. Tissues were infiltrated through graded acetone/EMBed 812 (Electron Microscopy Sciences, Hatfield, USA) mixtures (3:1, 1:1, 1:3 v/v) prior to final embedding in pure epoxy resin. Polymerization was achieved through sequential thermal curing (37 °C X 12 h → 60 °C X 48 h). Semithin sections (1 μm) were initially cut for toluidine blue screening. After positioning, the ultrathin sections (60-80 nm) were prepared using a diamond knife (Diatome, Hatfield, USA) on a Leica EM UC7 ultramicrotome (Leica Microsystems, Wetzlar, Germany), and the tissues were fished out onto the 150 meshes cuprum grids with formvar film (Servicebio, Wuhan, China). The sections were then sequentially stained with 2% uranyl acetate in ethanol and 2.6% lead citrate. Grid-mounted sections were examined using a Hitachi HT-7800 transmission electron microscope (Hitachi, Tokyo, Japan). Digital micrographs were acquired with an AMT XR-16 CCD camera system (Advanced Microscopy Techniques, Woburn, USA), with image analysis performed using ImageJ (NIH) and TEMography v3.2 software.

### PLA

PPIs in bEnd.3 cells or primary cerebrovascular ECs with or without gene modification were detected by Duolink In Situ Fluorescence PLA kits (Olink Bioscience, Uppsala, Sweden). The experimental procedure was according to manufactureŕs instruction with mild modification. In general, before starting PLA, bEnd.3 or primary cerebrovascular ECs were pretreated with or without experimental reagents. The pretreated or untreated samples were sequentially fixed with 4% PFA at room temperature for 15 min, permeabilized in 0.1% Triton X-100 PBS at room temperature for 5 min after washing twice (5 min X 2), and blocked in blocking solutiuon provided by the kit at 37°C for 30 min. The following incubation of primary antibodies and PLA probes, ligation and amplification, and mounting the slides for imaging were as described in product manual.

### Brain section IF staining and imaging

Mice to be tested were anesthesized and transcardially perfused with saline followed with 4% PFA for 4 h. The brains were then dissected and stored in 4% PFA at 4°C overnight. Then, paraffin sections or frozen sections were prepared as follows. For preparation of paraffin sections, the brains were dehydrated in increasing gradients of alcohol (70%-96%-100%, every change 30 min) and 100% Xylene (3 X 20 min). Next, the brains were put in freshly melted paraffin wax (58°C) for embedding and allowed to cool down overnight under room temperature, and 3 µm-thick slices were prepared with leica biosystems microtome (Leica Microsystems, Wetzlar, Germany). For preparation of frozen sections, the 4% PFA fixed brain tissues were cryoprotected in 30% sucrose until they sink. Then samples were embedded in OCT in cryomolds and snap frozen in prechilled isopentane or on dry ice. Frozen tissues can be stored in a -80 freezer and/or 10 µm-thick slices were prepared with leica biosystems cryostat (Leica Microsystems, Wetzlar, Germany). Before incubation with the first antibody, paraffin slices were deparaffinized and rehydrated by incubations with graded xylene and ethanol in water (Xylene 2 X 5 min, 1:1 Xylene:100% ethanol 5min, 100% ethanol 2 X 5min, 95% ethanol 5min, 70% ethanol 5 min, 50% ethanol 5 min, Deionized water 2 X 5 min). Then, the following steps were used for both paraffin and frozen sections. Slices were sequentially baked in a 37 ℃ oven for 10-20 min, fixed in 4% PFA for 30 min, incubated with 1mM EDTA antigen repair buffer (pH 8.0) for appropriate time, permeabilized with phosphate-buffered saline (PBS) containing 0.4% Triton X-100 and 1% donkey serum for 20 min, and blocked with PBS containing 0.1% Triton X-100 and 10% donkey serum or 3% BSA for 1 hour. Afterwards, the tissue slices were incubated with first antibody overnight at 4 ° C in a wet box, and after washing thrice incubated with appropriate fluorescent secondary antibodies at room temperature in dark for 1 h. The following sealing and mounting of the slides for imaging are as described in standard protocol. The images were captured using Pannoramic 250 Flash III or Pannoramic Midi and viewed with CaseViewer 2.4 (3DHISTECH, Budapest, Hungary) and analysed with ImageJ software.

### Cell IF staining and imaging

Cells were permeabilized with phosphate-buffered saline (PBS) containing 0.1% Triton X-100 and 1% donkey serum for 20 min, and blocked with PBS containing 0.1% Triton X-100 and 10% donkey serum or 3% BSA for 1 hour. Afterwards, the cells were incubated with first antibody at room temperature for 1 h in a wet box, and after washing thrice incubated with appropriate fluorescent secondary antibodies at room temperature in dark for 1 h. The following sealing and mounting of the slides for imaging are as described in standard protocol. The images were captured using confocal microscope Leica TCS SP8 X using 488 nm and 594 nm excitation lasers (Leica Microsystems, Wetzlar, Germany). All images acquired were analysed using LAS X software (Leica Microsystems, Wetzlar, Germany).

### Masson, EVG and ALP staining and imaging

Paraffin-embedded sections of mouse brain tissue from the experimental group were prepared and subsequently deparaffinized following standard protocols. All dyes used were from Servicebio (Wuhan, China) and the staining protocols were carried out according to manufacturer’s instruction. Slides were cleared with anhydrous ethanol and xylene, and then sealed with neutral gum. Images were acquired using Panoramic 250 Flash III or Panoramic Midi II and viewed with CaseViewer 2.4 (3DHISTECH, Budapest, Hungary).

### Co-IP

5x10^6^ cells or 10 mg homogenized brain tissue were suspended in 600 μL of lysis buffer (50 mM Tris-HCl pH 7.4, 1% Triton X-100, 1% n-Octyl β-D-glucopyranoside, 0.1% SDS, 1mM EDTA pH 7.0, 150 mM NaCl) supplemented with 1 X protease inhibitor cocktail (MERCK, Darmstadt, Germany) and incubated for 1 h on ice. Then, the lysates were centrifuged at 12,000 rpm for 10 min at 4°C to get rid of cellular nuclei and debris. The supernatant was transferred to fresh tube and protein concentration was measured using Pierce BCA Protein Assay Kit (Thermo Fisher Scientific, Waltham, USA). According to acquired protein concentration, appropriate amount of protein G agarose beads (MERCK, Darmstadt, Germany) resuspended in lysis buffer was added to preclear the non-specific bindings at 4°C for 30 min in rotating state. Meanwhile, target protein antibody conjugated beads or isotype antibody conjugated control beads were prepared by routine incubation and centrifugation protocol. Finally, the precleared supernatants and target protein antibody conjugated beads or isotype antibody conjugated control beads were combined and incubated for 2 h at 4°C in mild agitation state to pull down interested proteins and its interaction partners. After thrice centrifugation and washing with lysis buffer, the beads were resuspended in 100 μL lysis buffer and for each immunoblot 20 μL was used.

### scRNA-seq of mouse hippocampus

Single-cell suspensions were prepared from hippocampal tissues pooled from three mice per experimental group. Following quality assessment, cellular suspensions were processed through the Chromium Controller (10X Genomics, Pleasanton, USA) to generate single-cell Gel Bead-In-Emulsions (GEMs) using Chromium Next GEM Single Cell 3’ Reagent Kits v3.1 (10X Genomics, Pleasanton, USA). Each GEM compartment contained a hydrogel bead releasing oligonucleotides comprising an Illumina R1 sequencing primer (Read 1), a 16-nucleotide 10x cellular barcode, a 10-nucleotide unique molecular identifier (UMI), and a poly-dT primer for mRNA capture. Reverse transcription was performed within GEMs to synthesize barcoded cDNA. Post-GEM cleanup was conducted using silane magnetic beads (Thermo Fisher, Waltham, USA) to remove residual biochemical reagents. Full-length cDNAs were amplified and Library was constructed via sequential enzymatic processing. The final libraries contained the P5 and P7 primers used in Illumina bridge amplification. High-throughput sequencing on the final libraries was performed using Illumina NovaSeq X Plus (Illumina, San Diego, CA). Read 1 and Read 2 were standard Illumina® sequencing primer sites used in paired-end sequencing. Initial data processing was performed using Cell Ranger (v7.1.0, 10X Genomics) with the following quality thresholds: valid barcode rate >95% across all samples, median genes per cell ≥1,500, mitochondrial gene content <20%. Low-quality cells (<200 genes/cell or >5% mitochondrial reads) were excluded prior to downstream analysis. scRNA-seq datasets were harmonized using Harmony (v0.1.1). Top 30 principal components (PCs) explaining >85% variance were selected. Graph-based clustering was implemented through Scanpy (v1.9.3) using Leiden algorithm (resolution=0.8). Cell identities were determined by marker gene expression with thresholds (avg_log2FC >1.0, p_val_adj <0.01), cross-referencing with brain RNA-Seq database (v2022.1), and further by manual validation using canonical markers. UMAP projection was computed using Seurat (v4.3.0) with 30 neighbors and 0.3 minimum distance differential expression. The significantly up- and downregulated genes identified across distinct cellular subpopulations were systematically analyzed through Gene Ontology (GO) term enrichment and KEGG pathway mapping to uncover their predominant biological functions and molecular network associations. Furthermore, comparative functional profiling of intergroup differentially expressed genes (DEGs) was performed using comprehensive GO enrichment and KEGG pathway analyses, enabling precise characterization of pathway activation states and mechanistic insights into the biological processes underlying phenotypic variations between experimental groups.

### Cellular fractionation

Minute^TM^ plasma membrane raft isolation kit (invent biotechnologies, Beijing, China), peroxisome isolation kit (Sigma-Aldrich, Saint Louis, USA), endoplasmic reticulum (ER) isolation kit (Novus biologicals, Saint, Louis), cytoplasmic and nuclear extraction kit and mitochondria isolation kit (Beyotime, Shanghai, China) were used for cellular fractionation according to manufacturer’s instruction. Unless otherwise specified, high-speed centrifugation was performed using an Optima XPN-100 Ultracentrifuge (Beckman Coulter, Brea, USA).

### Immunoblot

Cells or homogenized brain tissue were suspended in appropriate volume of RIPA lysis buffer (50 mM Tris-HCl pH 7.4, 1% Triton X-100, 1% sodium deoxycholate, 0.1% SDS, 1mM EDTA pH 7.0, 150 mM NaCl) supplemented with 1 X protease inhibitor cocktail (MERCK, Darmstadt, Germany) and with or without 1 X phosphatase inhibitor cocktail (MERCK, Darmstadt, Germany) and incubated for 1 h on ice. Then, the lysates were centrifuged at 12,000 rpm for 10 min at 4°C to get rid of cellular debris. The supernatant was transferred to fresh tube and protein concentration was measured using Pierce BCA Protein Assay Kit (Thermo Fisher Scientific, Waltham, USA). Then, 30 μg cell or tissue lysate or 20μL immunoprecipitate for each sample was mixed with equal volume of 2 X loading buffer (4% SDS, 10% 2-Mercaptoethanol, 20% Glycerol, 0.004% Bromophenol blue, 0.125 M Tris-HCl, pH 6.8) and heated at 95°C for 10 min. The following Sodium Dodecyl Sulfate-Poly Acrylamide Gel Electrophoresis (SDS-PAGE) and Western blotting were according standard protocol. The blotted polyvinylidene fluoride (PVDF) or nitrocellulose (NC) membrane was blocked with 3% Bovine serum albumin (BSA) at room temperature (RT) for 1 h. Then, first antibody incubation (30 min to 1 h), Phosphate-buffered saline, 0.1% Tween 20 (PBST) or Tris-buffered saline, 0.1% Tween 20 (TBST) washing (3 X 10 min), and Horseradish peroxidase (HRP)-labelled secondary antibody incubation (1 h) were sequentially carried out at RT. At last, the membrane was incubated with enhanced chemiluminescence (ECL) chromogenic substrate solution for 1 min at RT, and the signals were recorded with film or charge-coupled device (CCD) cameras according to manufacturer’s instruction and analysed with ImageJ software.

### Isolation and culture of primary cerebrovascular ECs and brain astrocytes

To isolate primary ECs from experimental mice, cerebral microvessels were first prepared following a previously described protocol^88^. In brief, mice were anesthetized in a CO_2_ chamber and sacrificed by cervical dislocation, after which brains were removed and transferred to a petri dish with MCDB 131 medium (Thermo Fisher Scientific, Waltham, USA). The cortical and hippocampal tissues were homogenized in a Dounce tissue grinder and centrifuged at 2,000 × g for 5 min at 4°C. The pellets were resuspended in 15% dextran (in DPBS) and centrifuged at 10,000 × g for 15 min at 4 °C. After resuspending the resulting pellets in an appropriate volume of DPBS and filtering through a 40-µm cell strainer, microvessels were retrieved using MCDB 131 medium supplemented with 0.5% fatty acid-free BSA. (Beyotime Biotechnology, Shanghai, China). The extracted microvessels were then digested with 0.5 mg/ml collagenase (Sigma-Aldrich, Saint Louis, USA) or Liberase TH (Roche, Basel, Switzerland) for 30 min at 37°C to generate a single-cell suspension. After filtering through a 40-µm cell strainer, ECs were isolated using CD31 MicroBeads (Miltenyi Biotec, Bergisch Gladbach, Germany) according to the manufacturer’s instructions and were seeded on gelatin (Sigma-Aldrich, Saint Louis, USA) coated plate or coverslip and cultured at 37 °C/ 5% CO₂ in MCDB 131 medium supplemented with 5% FBS, 0.5% fatty acid-free BSA, 20 ng/mL EGF (Abbkine, Wuhan, China), 10 ng/mL bFGF, 10 ng/mL IGF-1, 1 μg/mL hydrocortisone, 10 μg/mL transferrin, 50 μg/mL ascorbic acid, 10 μg/mL heparin (MCE, Shanghai, China), 100 U/mL penicillin, and 100 μg/mL streptomycin. Cerebral astrocytes were isolated using Anti-ACSA-2 MicroBeads Kit combined with CD11b MicroBeads (Miltenyi Biotec, Bergisch Gladbach, Germany) according to the manufacturer’s instructions.

### FDISCO imaging and three-dimentional reconstruction of the mouse hippocampal vascular system

To label mouse blood vessels, 100 μL of Alexa Fluor 647-conjugated anti-mouse CD31 antibody (Jarvisbio, Wuhan, China) was administered via tail vein injection. Following injection, animals were placed in a warm cage for 30 minutes. Animals were then anesthetized with pentobarbital sodium and transcardially perfused sequentially with phosphate-buffered saline (PBS) and 4% paraformaldehyde (PFA). The hippocampus was dissected, post-fixed overnight in 4% PFA at 4°C, and rinsed twice with PBS to remove residual fixative. Hippocampal tissues were made transparent using a FDISCO (Farnesol-X Disabled Solvent-based Clearing with Outstanding imaging depth) tissue clearing kit (Jarvisbio, Wuhan, China) according to the manufacturer’s instruction. Cleared samples were imaged using a light-sheet microscope (LiToneXL; Light Innovation Technology, Kitakyushu, Japan) equipped with a 10x objective (NA > 0.5, working distance = 5 mm). Samples were illuminated from four sides using thin light sheets, and images were acquired as merged Z-stacks. During imaging, cleared samples were manually secured onto a sample holder adapter and immersed in a 3D-printed chamber filled with imaging solution/reagent. For image processing, original TIFF image files were stitched and converted using LitScan software (Light Innovation Technology, Kowloon, Hong Kong). 3D reconstruction, image capture (screenshots), and video generation were performed using Imaris software (Bitplane/Oxford Instruments).

### Targeted metabolomics for steroid hormone profiling

An ultra-high performance liquid chromatography coupled to tandem mass spectrometry (UHPLC-MS/MS) system (ExionLC™ AD UHPLC-QTRAP® 6500+, SCIEX, Marlborough, USA) was used for steroid hormone profiling of TPP2 depleted and WT cerebrovascular ECs. All of the 38 steroid hormone standards and 4 stable isotope-labeled standards were obtained from ZZ Standards Co., LTD. (Shanghai, China). Methanol, acetonitrile, isopropanol and acetic acid (Optima LC-MS Grade) were purchased from Thermo Fisher Scientific (Waltham, USA). Ultrapure water was purchased from Millipore (Burlington, USA). The stock solution of individual steroid hormone was mixed and prepared in steroid-free matrix to obtain a series of steroid hormone calibrators at a concentration of 0.25-25000 ng/mL. Certain concentrations of Testosterone-D4 、 Cortisol-D4 、 Progesterone-D9 or Cholesterol-D7 were compounded and mixed as Internal Standard (IS). Separation was performed on a Kinetex C18 column (2.1 × 100 mm, 1.7 μm) maintained at 50 °C. The mobile phase consisted of 0.04% acetic acid in 30% acetonitrile (v/v) (solvent A) and acetonitrile/isopropanol (6:4, v/v) containing 0.04% acetic acid (solvent B), delivered at a flow rate of 0.30 mL/min. The following gradient program was used: initial 5% B, 6 min; 5-25% B, 10 min; 25-55% B, 25 min; 55-100% B, 26 min; 100% B, 26.1 min; 100-5% B, 28 min; 5% B for re-equilibration. The mass spectrometer was operated in positive/negative multiple reaction mode (MRM) mode. Parameters were as follows: IonSpray Voltage (4500V/-4500 V), Curtain Gas (35 psi), Ion Source Temp (550°C), Ion Source Gas of 1 and 2 (60 psi).

LC-MS was used to analyze the calibration standards. Linear regression analysis was performed using the ratio of the standard concentration to the internal standard concentration (x-axis) versus the ratio of the standard peak area to the internal standard peak area (y-axis). A correlation coefficient (r) > 0.99 for each metabolite was required. The limit of quantification (LOQ) was determined using the signal-to-noise ratio (S/N) method, comparing the signal from the lowest measurable standard concentration to that of a blank matrix. Typically, the concentration yielding an S/N ratio of 10:1 was defined as the LOQ. Matrix comprises all sample components other than the target analyte. Matrix components can significantly interfere with the target’s analysis, affecting result accuracy; these interferences are termed matrix effects. To assess the matrix effects of steroid hormones on analyte ionization, the following calculation was used: ME% = [(QC matrix - B blank) / QC MS water-1] × 100%. Here, ME is matrix effect, QC matrix is blank matrix with standard sample, QC MS water is matrix free with standard sample, and B blank is blank matrix sample. Matrix effects were classified as minimal (-20% ≤ ME% ≤ 20%), moderate (ME% ≤ -20% or ME% ≥ 20%), or significant (ME% ≤ -50% or ME% ≥ 50%). Accuracy, expressed as recovery (%), reflects the closeness of measured results to the reference value. The accuracy for this method can be calculated as follows: R% = (QC recovery sample - B blank) / S × 100%. R% is recovery rate, QC recovery sample is matrix with recovery point sample, B blank is blank matrix sample, S is theoretical concentration. The accuracy for this mehod was determined at three different concentration levels. The accuracy requirement 85% to 115% and RSD ≤ 15% of three different concentration levels. Stability was evaluated to assess the integrity of the target analyte both in the biological matrix and in the processed sample. This method specifically assessed post-preparative stability in the autosampler (4°C) for 24 hours. The acceptance criterion was RSD ≤ 15% for each analyte concentration over this period.

### Fluorescent labeling of choline and its uptake assay

Three distinct ways, differing by the conjugation site and fluorophore, were used for fluorescent labelling of choline. Of which, choline has either NBD fluorophore conjugated to the methyl group (Choline-CH3-NBD) or CY2 fluorophore conjugated to the hydroxyl group (Choline-O-CY2) or to the carbon atom adjacent to hydroxyl group (Choline-CO-CY2). The labelling protocols were carried out by Ruixi biotechnology (Xian, China) and Xinweichuang biology (Chongqing, China) respectively. The purity of all final products is above 95%, and their quality has been verified through mass spectrometry testing. The final product was dissolved in Dimethyl sulfoxide (DMSO) or methanol to prepare a 100mM stock solution. For the uptake assay, stock solution was diluted to 50 µM using phenol red free DMEM culture medium (Thermo Fisher Scientific, Waltham, USA) for cell incubation. For confocal microscopy imaging, cells were incubated at 37 °C and 5% CO₂ for 30 min, and then fixed in 4% PFA for 10 min. For dynamic analysis of choline uptake, live imaging was performed for 5 minutes using live cell station of Nikon AX with NASPARC confocal microscope system (Nikon, Tokyo, Japan).

### Assessment of cell membrane fluidity

Cell membrane fluidity TMA-DPH fluorescence detection kit (Bestbio, Shanghai, China) and BODIPY FL C5-HPC (MCE, Shanghai, China) were used according to manufacturer’s instruction. In brief, cells were seeded in BeyoGold ™ Black transparent 96 well cell culture plate (Beyotime, Shanghai, China) and washed once with pH 7.4 PBS buffer after growing to 80% confluency w/o drug treatment. Cells were then incubated at 37 °C for 30 min with 1:1000 diluted TMA-DPH probe prepared in the working solution provided in the kit or BODIPY FL C5-HPC solution. After washing once with pH 7.4 PBS, fluorescence polarization was measured at Ex/Em: 360/460 using BioTek Synergy H1 (Agilent Technologies, Santa Clara, USA) or VICTOR Nivo plate reader (PerkinElmer, Springfield, USA). Fluorescence polarization value (mP) was calculated as following formula: P = (S-G·P)/(S+G·P). In the formula S represents the emission light intensity (RFU) when the two polarizers are parallel to each other, P represents the RFU when the two polarizers are perpendicular to each other, and G is a correction factor of the measurement device.

### Protein-lipid molecular docking

Molecular docking experiments were performed using MOE2019 (Chemical Computing Group ULC, Montreal, Canada), during which a total of 30 conformations were collected. The binding pocket was predicted with MOE, and conformation sampling and scoring were conducted using the induced fit method. The first-round scoring function was London dG, and the second-round scoring function was GBVI/WSA dG. Based on the docking scores, the top 10 conformations with the highest scores were selected and further filtered according to the binding site. Interaction diagrams were generated using Discovery Studio software (Biovia, San Diego, USA), and binding diagrams were created with PyMOL(Schrödinger, New York, USA).

### AAV virus production and injection

The mouse Far1 cDNA was amplified by PCR using C57BL/6 mouse genomic DNA as templates, phanta super-fidelity DNA polymerase (Vazyme, Nanjing, PR China), and the following forward and reverse primers: 5’- CCACAGCAACTGACCCCGGGGGATCCGCCACCatggtttcaatcccagaat-3’ and 5’-GTAGTCGTTAATTAAGGTACCACTAGTgtatctcatagtgctggatg-3’. The acquired cDNA after sequencing validation was cloned into the adeno-associated virus type 2 (AAV2) derived vector pHBAAV-Tie1-MCS-Flag-ZsGreen (HANBIO, Shanghai, PR China) between BamHI and KpnI cutting sites to get the final plasmid pHBAAV2/Br1-Tie1-Far1-Flag-ZsGreen. The recombinant plasmid pHBAAV2/Br1-Tie1-Cyp19a1-Flag-T2A-ZsGreen was generated by subcloning the mouse Cyp19a1 cDNA from the donor vector pHBAAV2/BBB-hSyn-Cyp19a1-T2A-ZsGreen into the recipient vector pHBAAV2/Br1-Tie1-MCS-Flag-T2A-ZsGreen. After high purity endotoxin free extraction, the final plasmid was co-transfected with pAAV-RC and pHelper into AAV-293 cells using Lipofiter^TM^ transfection reagent (HANBIO, Shanghai, PR China). Cell precipitates were harvested 72 h after transfection and high titer virus preservation solution was obtained after column purification, and various indicators of virus were determined according to strict quality standards (HANBIO, Shanghai, PR China). Then, 50 μL (1 X 10^11^ vector genome (vg)) recombinant virus or control virus particles per mouse were delivered through injection into the tail vein. One month later, the learning and memory abilities of the experimental mice were measured using the MWM test. Expression of the recombinant Far1 or Cyp19a1 was validated by whole brain imaging using IVIS Lumina III in vivo imaging system (PerkinElmer, Watham, USA) and by immunoblot.

### Morris water maze test

The Morris Water Maze (MWM) test was performed following a published nature protocol ^89^ with minor modifications. Briefly, a circular pool (120 cm in diameter) with a hidden platform (10 cm²) placed in one quadrant was filled with tap water until the platform was submerged 1 cm beneath the water surface. The platform was then camouflaged by placing opacifying titanium dioxide in the water and the water temperature was maintained at 23°C. During the 5-day acquisition phase, each mouse underwent four trials per day. The starting position was varied across the four trials such that the platform was positioned to either the right or left of the animal at the start, with one trial initiated from each of the four cardinal starting points daily. If a mouse failed to locate the platform within 60 s, it was gently guided to it and allowed to remain there for 30 s. The time taken to find the platform was recorded as the escape latency. On day 6, a probe trial was conducted with the platform removed. Mouse behavior was recorded for 60 s using EthoVision XT video tracking system (Noldus, Leesburg, USA) to measure the time spent in the target quadrant, swimming speed, total distance moved, and the number of crossings over the former platform location, in order to evaluate spatial memory retention.

### Statistical analysis

All statistical analyses were performed using GraphPad Prism 10.1.2 (La Jolla, USA). Unless otherwise specified, data are expressed as mean ± SEM from at least six independent experiments, except for Western blot (WB) results, which were derived from three independent experiments. For comparisons between two groups, parametric data were analyzed using the two-tailed unpaired Student’s t-test, while nonparametric data were assessed with the Mann–Whitney U test. For comparisons among more than two groups, parametric data were evaluated by one-way ANOVA followed by Bonferroni’s post hoc test, and nonparametric data were analyzed using the Kruskal–Wallis test. The escape latencies across different mouse groups were compared using a two-way repeated measures analysis of variance (ANOVA), with post hoc comparisons performed via Tukey’s test. Normality of data distribution was verified using the Kolmogorov–Smirnov goodness-of-fit test.

## Supplementary material

Suppl. antibody table 1, Suppl. Fig. 1, Fig. 2 and Fig. 3

## ACKNOWLEDGMENTS

This work was supported by the intramural program of the Henan Province Mental and Neurological Disease Dominant Discipline Construction Project of Henan Medical University to J. H and J. Z (505308). Young Scientists Fund of the National Natural Science Foundation of China to X. Z (82401652). The content of this manuscript is solely the responsibility of the authors and does not represent the official views of the funding agency. We thank all members of Institute of Psychiatry and Neuroscience, Henan Medical University for providing overall support for implementing of this study. We extend our sincere gratitude to Professor Zhaobing Gao, Professor Liu Cao, Professor Weidong Wu, Professor Chengbiao Lu and Dr Jia Tong for their comprehensive and strong support. We thank Mr Xiaoyu Zhou for helping with PET/CT and NMR/MRI and Ms Zeqing Wu for helping with the whole brain ex vivo imaging.

## AUTHOR CONTRIBUTIONS

Conceptulization: J. Huai; J. Zhao. Methodology: J. Zhao; J. Huai; X. Zhang; J. Zhao; J. Liu; H, Zhang; R. Zhang; S. Yang. Investigation: J. Zhao; Q. Guo; X. Xin; Y. Zhang; K. Pei; J. Li; G. Wang; J. Li; W. Sun; S. Huang; X. Fan; L. Luo; W. Li; Y. Wang; Y. Cao; B. Yang. Writing and editing: J.Huai; J. Zhao. Project administration, J. Huai; J. Zhao.

## DELARATION OF INTERESTS

The authors declare no competing interests..

---

**Suppl. Fig. 1.**
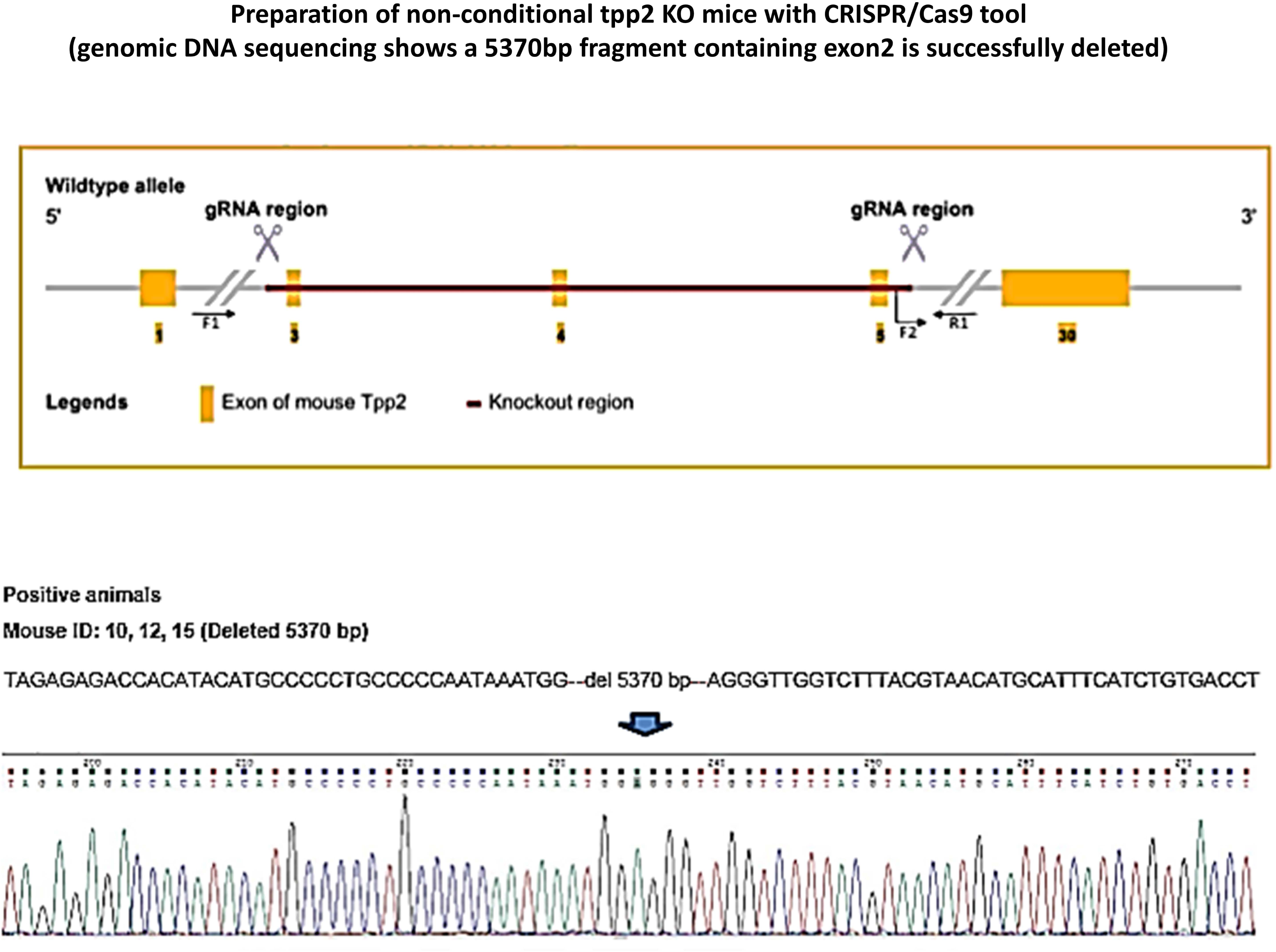

**Suppl. Fig. 2.**
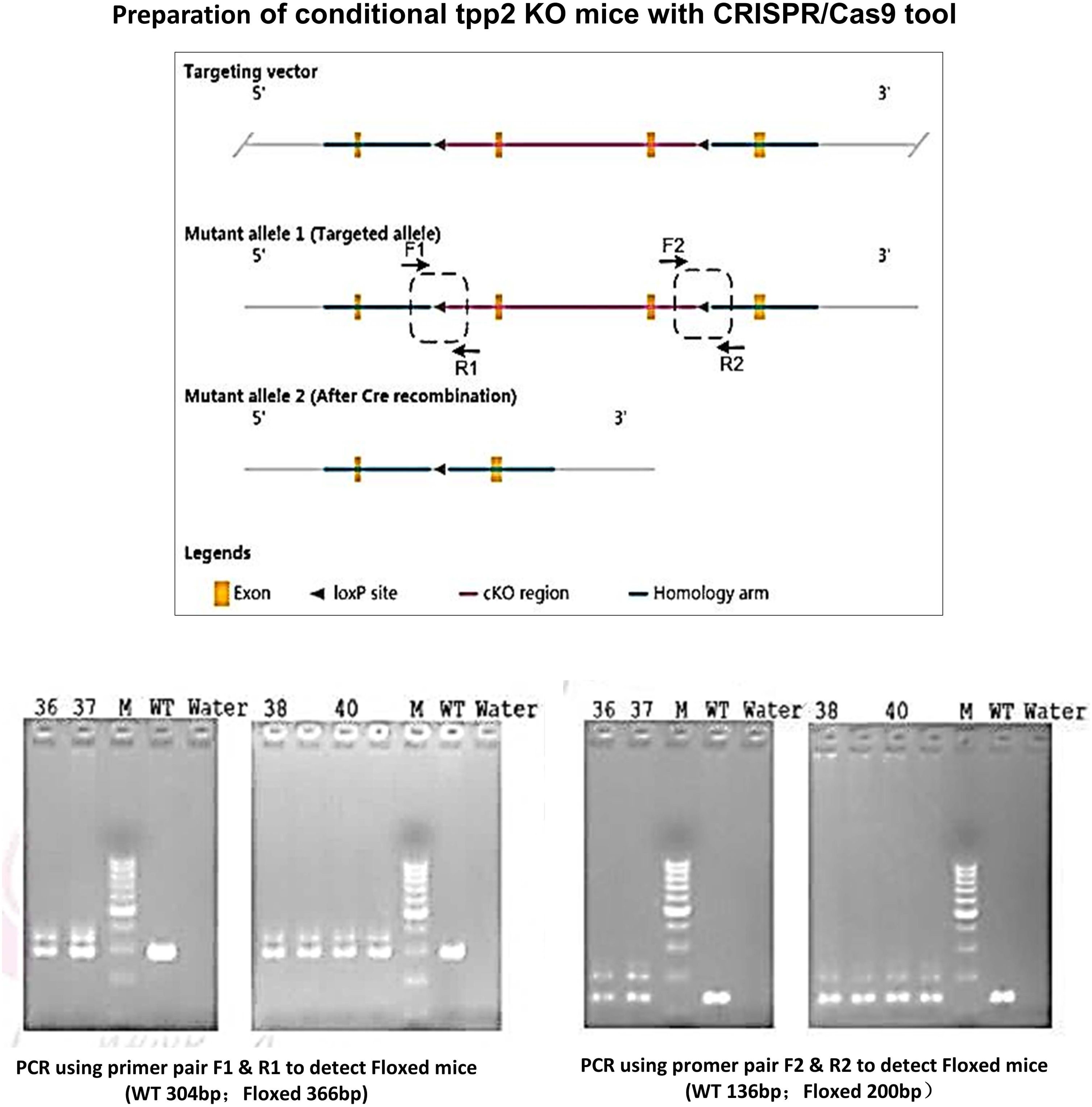

**Suppl. Fig. 3.**
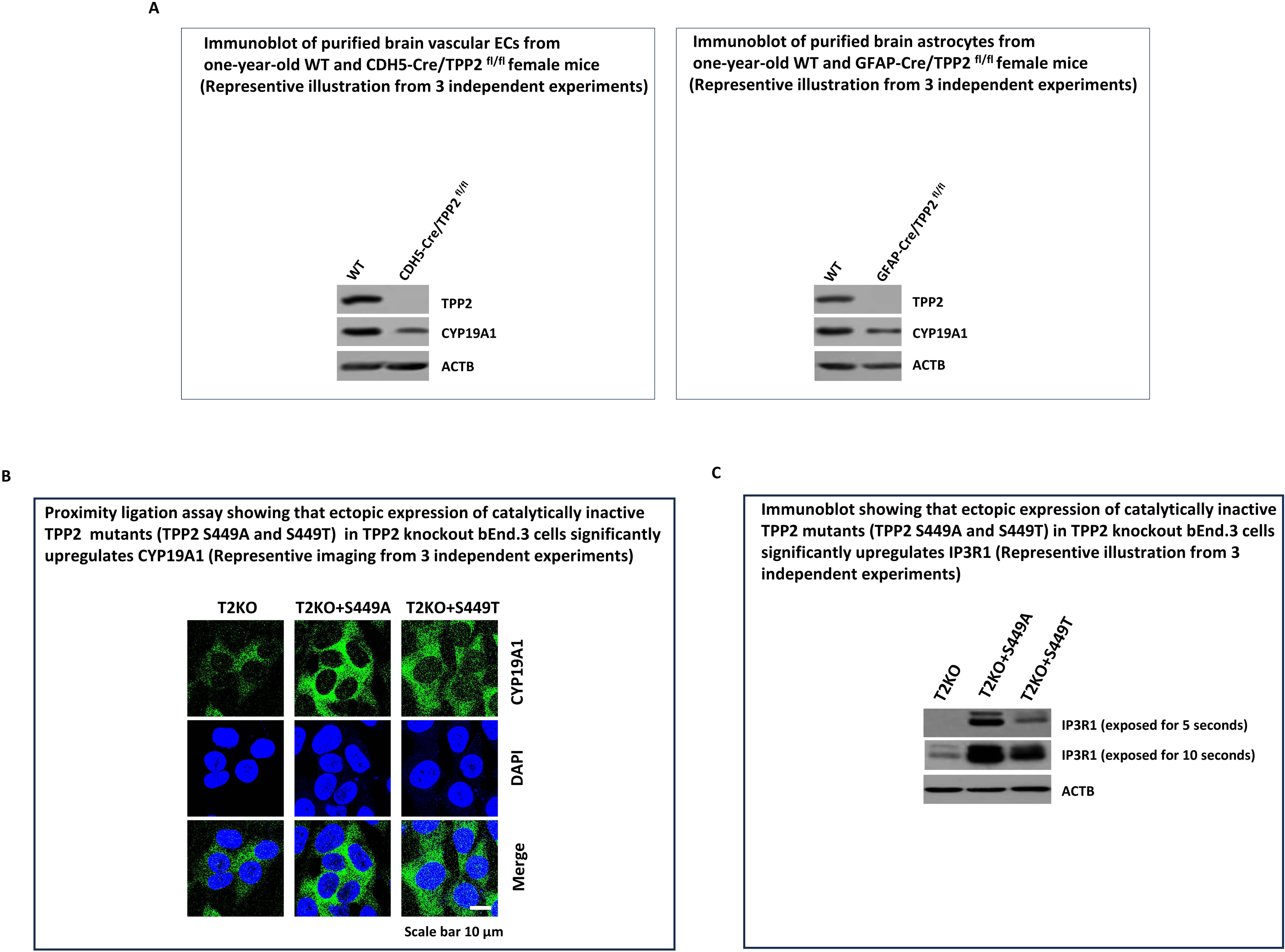

**Supplementary table 1.**

| Name | Antibodys | Source | Identifier |
| --- | --- | --- | --- |
| ACBD4 | ACBD4 Polyclonal antibody | Proteintech | 20941-1-AP |
| ACBD5 | ACBD5 Polyclonal antibody | Proteintech | 21080-1-AP |
| ACTB | β-Actin Rabbit mAb (High Dilution) | Abclonal | AC026 |
| AGPS | Alkyl-DHAP synthase/AGPS Polyclonal antibody | Proteintech | 21011-1-AP |
| ATF6 | ATF-6 (D4Z8V) Rabbit Monoclonal Antibody | Cell Signaling | 65880S |
| ATF6 | ATF6 monoclonal antibody | Proteintech | 66563-1-Ig |
| CALR | Calreticulin (D3E6) XP <sup>®</sup> Rabbit mAb | Cell Signaling | 12238S |
| CAM | Calmodulin (D1F7J) Rabbit mAb | Cell Signaling | 35944S |
| Camk2 | CaMKII Recombinant Rabbit Monoclonal Antibody [SU03-57] | HUABIO | ET1608-47 |
| Catalase | Catalase Monoclonal antibody | Proteintech | 66765-1-Ig |
| CD31 | Anti-CD31 Mouse mAb | Servicebio | GB120005 |
| CHIP | CHIP (C3B6) Rabbit Monoclonal Antibody | Cell Signaling | 2080S |
| COMT | COMT Rabbit Polyclonal Antibody | Proteintech | 14754-1-AP |
| CYP19A1 | Aromatase Antibody | Novus | NB100-1596 |
| CYP19A1 | Rabbit anti-Aromatase Monoclonal Antibody(3D8) | Absin | abs159190 |
| ERα | Estrogen Receptor alpha Monoclonal Antibody (6F11) | Invitrogen | MA1-80216 |
| FAR1 | Rabbit Polyclonal FAR1 antibody | Novus | NBP1-89847 |
| FLAG | Flag-Tag Mouse Monoclonal Antibody | Abways | AB0008 |
| GNPAT | GNPAT Polyclonal antibody | Proteintech | 14931-1-AP |
| IP3R1 | IP3 Receptor 1 (D53A5) Rabbit mAb | Cell Signaling | 8568S |
| IRE1 | IRE1 alpha (14C10) Rabbit Monoclonal Antibody | Cell Signaling | 3294S |
| MARCH5 | MARCH5 Antibody | Cell Signaling | 19168S |
| p62 | BD Transduction Laboratories™ Purified Mouse Anti-Nucleoporin p62 | BD Biosciences | 610497 |
| PERK | PERK (C33E10) Rabbit Monoclonal Antibody | Cell Signaling | 3192S |
| PEX14 | PEX14 Polyclonal antibody | Proteintech | 10594-1-AP |
| PEX16 | PEX16 Monoclonal antibody | Proteintech | 68261-1-Ig |
| PEX19 | PEX19 Polyclonal antibody | Proteintech | 14713-1-AP |
| PEX2 | PXMP3 Antibody | Novus | NBP2-94643 |
| PEX3 | PEX3 Polyclonal antibody | Proteintech | 30424-1-AP |
| PEX5 | PEX5 (D7V4D) Rabbit mAb | Cell Signaling | 83020S |
| PEX5 | PEX5 Antibody | Abmart | MG511430 |
| PEX7 | PEX7 Polyclonal antibody | Proteintech | 20614-1-AP |
| PIP2 | Anti-PIP2 antibody [2C11] | Abcam | ab11039 |
| PLB | Phospholamban Monoclonal Antibody (2D12) | Thermo Fisher | MA3-922 |
| PLCγ | PLC gamma 1 Antibody | Abmart | T55638 |
| PMP70 | PMP70 antibody | sigma | SAB4200181 |
| PMP70 | PMP70 Antibody | MCE | HY-P81883 |
| PMP70 | ABCD3 Polyclonal antibody | Proteintech | 26720-1-AP |
| SERCA1 | ATP2A1/SERCA1 (L24) Antibody | Cell Signaling | 4219 |
| SIGMAR1 | SIGMAR1 Polyclonal antibody | Proteintech | 15168-1-AP |
| SRC | c-SRC Polyclonal antibody | Proteintech | 11097-1-AP |
| SRC | c-SRC Monoclonal antibody | Proteintech | 60315-1-Ig |
| TPP2 | TPP2 Polyclonal antibody | Proteintech | 14120-1-AP |
| VDAC2 | VDAC2 Mouse Monoclonal Antibody | Proteintech | 66388-1-IG |
| α-SMA | Anti-alpha smooth muscle Actin Rabbit pAb | Servicebio | GB111364 |

